# State-Threatened Gopher Tortoise (*Gopherus polyphemus*) Gut Microbiome Analysis Reveals Health Insights into Southeastern Florida Population

**DOI:** 10.1101/2024.08.28.609265

**Authors:** Dimitrios S. Giakoumas, Jon Moore, Ericca Stamper, Kelsie M. Bernot, Tracy J. Mincer

**Affiliations:** Harriet L. Wilkes Honors College, Florida Atlantic University, Jupiter, FL, United States of America; Harbor Branch Oceanographic Institute, Florida Atlantic University, Fort Pierce, FL, United States of America

**Keywords:** Gopher Tortoise_1_, *Gopherus polyphemus*_2_, Gut Microbiome_3_, Abacoa Greenway_4_, Florida_5_

## Abstract

The gopher tortoise (*Gopherus polyphemus*), endemic to the southeastern part of the United States, is a keystone species that provides ecological support to over 350 species. Deforestation, urbanization, road mortality, and disease, over the past 100 years, have caused population decline by over 80%, causing Florida to declare the gopher tortoise a threatened species. Little is known about the gopher tortoise gut microbiome, which could play a major role in tortoise health. This study aimed to better understand the health and biology of this threatened keystone species through the characterization and analysis of their gut microbiomes in the Abacoa Greenway of Southeastern Florida using next-generation sequencing to survey the gut microbiome. Major findings include: high levels of alpha diversity; lack of significant difference of male and female alpha diversity and taxonomic composition profiles and, counterintuitively, male beta diversity was less disparate than female beta diversity; phyla associated with fiber fermentation short-chain fatty acid metabolism, Firmicutes and Bacteroidetes, predominated all samples. Notable probiotic taxa present included: Lachnospiraceae and *Clostridium butyricum*. Potentially pathogenic taxa included: *Helicobacter* sp. and Mollicutes sp. Pathogenetic taxa undetected: gopher tortoise *Helicobacter*, and *Mycoplasma* spp. These results add to the understudied reptiles and tortoise gut microbiome.

## Introduction

Next generation sequencing (NGS) technology has contributed to unearthing the complex and diverse functions and identities of unculturable microbes within their respective niches (Nam et al., 2023). In particular, much attention has been paid to the “gut microbiome”. The gut microbiome, which can be comprised of archaea, bacteria, protists, fungi, and viruses, has proved to play a critical, symbiotic role to the health of eumetazoans (Afzaal et al., 2022; Ling et al., 2022; Riehl et al., 2024; Salvadori and Rosso, 2024). Indeed, this enteric symbiosis has likely endured across the evolutionary history of eumetazoans, from *Hydra* (Dupre and Yuste, 2017) to *Homo sapiens*.

Microbe-host connections can be highly interwoven. For example, human and murine models show intricate neurological and immunological relationships with gut microbes. In fact, 70% of immune cells, i.e., the gut-associated lymphoid tissue, are associated with the gut (Rhee et al., 2004; Vighi et al., 2008; Ruth and Field, 2013; Wiertsema et al., 2021). Short chain fatty acids (SCFA) such as acetate, propionate, and butyrate, which are indigestible to vertebrates, are metabolized from polysaccharides by gut bacteria (Bedford and Gong, 2018; Liu et al., 2018). Butyrate has been shown to interact with gut-associated lymphoid tissue via histone deacetylation inhibition (Chang et al., 2014); additionally, it has been shown to improve the epithelial integrity in the gut and the blood-brain barrier, thereby preventing pathogenic penetration (Braniste et al., 2014), and regulating the transport of particles between a host’s internal and exterior milieus. Bacterial metabolites as such contribute to the metabolic up-regulation of 5-HT in enterochromaffin cells (Macfarlane and Macfarlane, 2011; Liu et al., 2021). Additionally, butyrate can provide up to 70% of energy to the colon (Clausen and Mortensen, 1995), maintaining the mucosal layer, and thereby prevents colonization and adhesion to the colonic epithelium of pathogenic endospores (Macfarlane and Macfarlane, 2011; Buffie and Pamer, 2013; Gonçalves and Martel, 2013). Additional examples of mechanisms for pathogenic immune defense include competition for resources, promotion of probiotics within the host (Zhu et al., 2023), and production of bacteriocins (Thomas et al., 2011; Buffie and Pamer, 2013; Heilbronner et al., 2021; Anjana and Tiwari, 2022). Specific probiotics may be used to fight disease; their metabolites may prevent endoparasites, thus yielding novel therapies (Saracino et al., 2021).

Categorized within the 2% of herbivorous reptiles (Vitt, 2004), the tortoise gut microbiome is only beginning to be described. However, despite its symbiotic relationship to disease, detailed attention to its relationship to health with disease in tortoises is lacking. Tortoises, being hind-gut fermenting, reptiles, and hence ectotherms, must facilitate fermentation year-round (Schaffner, 2015). Most of their diet thereby consists of fermentable yet indigestible, plant-derived starches, such as cellulose and hemicellulose. These starches are metabolized by gut bacteria and the metabolites are converted into energy and heat by their host.

Despite the significance of an animal’s gut microbiome to its health, the reptile gut microbiome has been understudied. Even more, the gut microbiome of the gopher tortoise (*Gopherus polyphemus*) has been understudied. To date, only three studies have provided insight into the gopher tortoise’s gut microbial makeup; two (Gaillard, 2014; Yuan et al., 2015) of which did not investigate populations in southeast Florida; one of which is an undergraduate thesis (Giakoumas, in press).

Gopher tortoises are burrowing land turtles found in the southeastern U.S. from Louisiana to South Carolina and throughout most of Florida (Bramble et al., 2014). They have an outsized importance in flatwoods, sandhills, high pine, scrub, and other ecosystems composed of well-drained sands. They are ecosystem engineers due to construction of burrows (Kinlaw and Grasmueck, 2012), which come to host an array of invertebrate and vertebrate commensal organisms, increasing biodiversity in the local area, thereby making gopher tortoises also a keystone species (Hansen, 1963; Eisenberg, 1983; Catano and Stout, 2015; Hipps, 2019). An additional element of gopher tortoises as keystone species is their zoochory or dispersal of seeds across local landscapes (Carlson et al., 2003; Hanish et al., 2020). Gopher tortoises are considered threatened in the state of Florida (Florida endangered and threatened species list; prohibitions, 2023). Enteric endoparasites and upper respiratory tract infections contribute to gopher tortoise population decline in Florida (Fremont, 2017; Huffman et al., 2018; Cooney et al., 2019; Page-Karjian et al., 2021).

Given the utility of metagenomic sequencing to uncover potential gut probiotics and pathogens and their relevance to immunity and health, we investigated, characterized, and analyzed the gopher tortoise gut microbiome of the Abacoa Greenway Range VIa. Our efforts aim contribute to the conservation of this vital species. This study builds upon Giakoumas’ (in press) undergraduate thesis.

## Materials and Methods

### Site Description

The Abacoa Greenway is a linear nature preserve segmented by multiple roadways crossing the area. The tortoises examined in this study were in Range VIa of the Abacoa Greenway in Jupiter, Florida (26.90°N, 80.11°W). The site is a mesic to xeric pine flatwoods ecosystem featuring about 9.27 hectares (22.9 acres) of more xeric upland dominated by slash pine (*Pinus ellioti*) and scrub oaks (*Quercus* spp.) and another 4.18 hectares (10.3 acres) of depressed mesic catchment basin for flood control dominated by slash pine, muhly grass (*Muhlenbergia capillaris*), and bald cypress (*Taxodium distichum*) (Hanish et al., 2020). The site is overcrowded containing more than 110 tortoises (i.e., 11.87 tortoises per hectare, Moore personal observation). The site and its vegetation are described further in Wetterer & Moore (2005) and Hanish et al. (2020).

### Sample Collection

All capture, handling, and sampling procedures were approved by the following: Florida Atlantic University IACUC approvals (A-19-41; A-22-47; A(T)22-06) and Florida Fish and Wildlife Conservation Permit (LSSC-14-00066B & C). Fecal samples (*n* = 26) from tortoises (*n* = 23) were acquired from the Abacoa Greenway Range VIa located in Jupiter, Florida between June 2022 and March 2023. Samples were immediately and nondestructively collected with sterilized forceps while tortoises were supinated and undergoing a physical examination. Each fecal sample was placed in a Ziplock bag, placed on a towel in an ice cooler, and transferred to a −80°C freezer in under four hours until processed.

### Target Description and Population

Gopher tortoises show a latitudinal gradient to age of maturity, which is related to growth rates. Germano (1994) noted that gopher tortoises across much of the species’ range mature at about 230 mm carapace length (CL), although several other authors have observed that females mature at 250 mm CL. More northerly populations in Georgia and Alabama grow slower and will sexually mature at 19-21 years old (Landers et al., 1982; Aresco and Guyer, 1999). Populations in northern Florida mature at 14-18 years (Berish and Leone, 2014). Doonan (1986) found a population in the Orlando area matured at 10-13 years of age. McLaughlin (1990) found faster growth rates with individuals achieving 230 mm CL by 8-9 years old and earlier maturity at 9-13 years old on Sanibel Island in southwestern Florida. Tortoises in southeastern Florida mature at 7-10 years of age and at that 230 mm CL threshold (Sano, 2014; Jon Moore personal observations, 2023).

Tortoises with a comparatively longer tail, longer gular projection, more concave plastron, or who were observed copulating were deemed to be adult males (McRae, Landers and Cleveland, 1981; Jon Moore, personal communication, 2023). Age was ascertained by counting the number and width of the growth rings embedded within each tortoise’s carapace scales (Germano, 1998; Jon Moore, personal communication, 2023); thin rings indicate sexual maturity, as energy is diverted away from growth to reproduction (Wilson et al., 2003). Just about all of the individuals at this site have been measured, aged, weighed, and permanently marked within the last 23 years and most of the individuals in this study have a long-known history of capture and recapture (Jon Moore, personal communication, 2023). Only four tortoises in this study represent likely waifs recently added to the site (T195, T203, T212, T216).

### DNA Extraction

Fecal samples were taken out of the −80°C freezer and thawed in their respective Ziplock bags for no more than 60 minutes. We extracted 100-200 µg of fecal and plant matter from the core of each fecal sample with tweezers, which were sterilized with ethanol between aliquots. DNA was extracted from (*n* = 26) fecal samples with the ZymoBIOMICS^TM^ DNA Miniprep Kit and the Quick-DNA^TM^ Fecal/Soil Microbe DNA MiniPrep Kit (cat. D4300 & D6010, respectively; Zymo Research, Irvine, CA, USA), in accordance with their respective protocols. Cell lysis was performed with ZR BashingBead Lysis Tubes using Zymo’s recommendations for the Vortex-Genie® 2, 120V: 40 minutes at the highest setting, “10”, with tubes taped horizontally on the rubber surface.

Purity and concentration for each DNA sample were measured using a Thermo Scientific^TM^ NanoDrop^TM^ 2000c Spectrophotometer (Thermo Fisher Scientific; Wilmington, United States). In accordance with Zymo Research’s downstream quality control standards, we achieved a 260/280 ratio ≥ 1.8 and surpassed or approximated a 230/260 ratio of ≥ 2.0.

### ZymoBIOMICS® Targeted Sequencing Service and Bioinformatics Analysis

Samples were then outsourced to Zymo Research Corporation, Irvine, CA, USA, for high-throughput, next-generation metagenomic sequencing (NGS) and bioinformatic analysis. The following methods have been modified from a generic materials and methods section from our ZymoBIOMICS**®** report:

### Targeted Library Preparation

Bacterial 16S ribosomal RNA gene targeted sequencing was performed using the *Quick*-16S^TM^ NGS Library Prep Kit (Zymo Research, Irvine, CA). The bacterial 16S primers amplified the V3-V4 region of the 16S rRNA gene. These primers have been custom designed by Zymo Research to provide the best coverage of the 16S gene while maintaining high sensitivity.

The sequencing library was prepared using an innovative library preparation process in which PCR reactions were performed in real-time PCR machines to control cycles and therefore limit PCR chimera formation. The final PCR products were quantified with qPCR fluorescence readings and pooled together based on equal molarity. The final pooled library was cleaned with the Select-a-Size DNA Clean & Concentrator^TM^ (Zymo Research, Irvine, CA), then quantified with TapeStation® (Agilent Technologies, Santa Clara, CA) and Qubit® (Thermo Fisher Scientific, Waltham, WA).

### Control Samples

The ZymoBIOMICS® Microbial Community Standard (Zymo Research, Irvine, CA) was used as a positive control for each DNA extraction, if performed. The ZymoBIOMICS® Microbial Community DNA Standard (Zymo Research, Irvine, CA) was used as a positive control for each targeted library preparation. Negative controls (i.e., blank extraction control, blank library preparation control) were included to assess the level of bioburden carried by the wet-lab process.

### Sequencing

The final library was sequenced on Illumina® MiSeq^TM^ with a v3 reagent kit (600 cycles). The sequencing was performed with 10% PhiX spike-in.

### Bioinformatics Analysis

Unique amplicon sequences variants were inferred from raw reads using the DADA2 pipeline (Callahan et al., 2016). Potential sequencing errors and chimeric sequences were also removed with the DADA2 pipeline. Taxonomy assignment was performed using UCLUST from Qiime v.1.9.1 with the Zymo Research Database, a 16S database that is internally designed and curated, as reference. Composition, alpha-diversity, and beta-diversity analyses were performed with Qiime v.1.9.1 (Caporaso et al., 2010). Analyses such as heatmaps, Taxa2ASV Decomposer, and PCoA were performed with internal scripts.

### Data Visualization and Statistical Analysis

Unless otherwise stated, all data visualizations and statistical analyses were performed using Python v.3.9. Outside of Qiime, analyses and plotting were carried out within the DataSpell IDE using Jupyter Notebook, R console, and R Markdown.

### Rarefaction Analysis

Rarefaction curves were generated using Qiime v.1.9.1 to assess the diversity of samples at different sequencing depths. The plots were generated using the Pandas v.1.5.3, plotly.graph_objects v.5.18.0, and Seaborn v.0.13.2 libraries (Plotly Technologies Inc. Collaborative data science., 2015; Waskom, 2021; pandas development team, 2023), with color maps provided by matplotlib.colors v.3.7.0 (Caswell et al., 2023).

### Relative Abundance Plotting

Relative abundance of different taxa was visualized using bar graphs. These graphs were generated using the Pandas v.1.5.3 and plotly.express v.5.18.0 libraries (Plotly Technologies Inc. Collaborative data science., 2015; pandas development team, 2023).

### Alpha Diversity Plots

Alpha diversity was visualized using GraphPad Prism v.10.2.3 (for Mac, GraphPad Software, Boston, Massachusetts USA, www.graphpad.com).

### Beta Diversity Analysis

To assess dispersion and variance in OTU Bray-Curtis beta diversity differences between males and females, as well as male and female samples, we performed PERMDISP and PERMANOVA tests. These analyses were conducted using the scikit-bio (skbio) library v.0.5.9 (Rideout et al., 2023) and the NumPy library v.1.24.2 (Harris et al., 2020) with 999 permutations. Since we obtained three temporally independent samples from a single female, T85 (i.e., T85.1, 2, and 3), we created four separate Bray-Curtis distance matrices using Qiime2 v.2019.7, including only one T85 sample (T85.1, 2, then 3) per analysis, and performed one analysis including all three samples to see if it made a difference in male-female PERMDISP and PERMANOVA results. Violin plots of Bray-Curtis beta diversity were visualized using GraphPad Prism v.10.2.3.

### Biplot and PCoA Analysis

The biplot analysis was performed using Qiime2. To visualize the relationship between variables, loadings, and samples was done using the matplotlib library v.3.7.0 (Caswell et al., 2023). The biplot provides a graphical representation of Bray-Curtis composition dissimilarity between samples and their feature loadings, which explain the dispersion of samples across each principal coordinate axis. The only sample that was excluded was from a juvenile.

Principal Coordinates Analysis (PCoA) was performed to further explore the compositional differences among the samples. The analysis included all samples including one from a juvenile (T215) that has not been sexed. The PCoA results were visualized using the matplotlib library v.3.7.0 (Caswell et al., 2023), and the principal coordinates were plotted to show the relative distances and clustering of the samples based on Bray-Curtis dissimilarity.

### Heatmap Generation

To visualize the differential taxa and ASV abundance at the species level between male and female samples, we generated heatmaps using the *pheatmap* package in R v.1.0.12 (Kolde, 2019).

### Differential Abundance: LEfSe Analysis

A linear discriminant analysis (LDA) effect size (LEfSe) (Segata et al., 2011) was used (LEfSe v.1.1.2, Python v.3.11.7; http://github.com/SegataLab/lefse/) to identify statistically significant taxa associated between male and female samples. LEfSe implements a series of statistical tests: first, a non-parametric factorial Kruskal–Wallis rank sum test; second, a pairwise test using the unpaired Wilcoxon sum-rank test; and finally, LDA to estimate the effect size of each differentially abundant OTUs. The analysis was conducted using default parameters except for the following: Logarithmic LDA score threshold for discriminative features: 2.5.

### Potential Pathogen and Probiotic Investigation

Investigation of potential pathogens and probiotics was conducted by searching through OTU composition table files for known gopher tortoise pathogens and probiotics.

### Post-Hoc Taxa Investigation and Analyses

Once taxonomic composition was obtained, we researched potentially relevant taxa that may be pathogenic or probiotic in gopher tortoises and other vertebrates. We performed Basic local Alignment Search Tool (BLAST) queries in the National Center for Biotechnology Information (NCBI) database (Zhang et al., 2000) for potential pathogens and probiotics as well. We calculated phylogenetic relationships via the online CLUSTAL Omega v.1.2.4 algorithm (Madeira et al., 2024); template scripts were generated using ChatGPT-4. To determine if the pathogenetic taxon *Mycoplasma* spp. could be present in tortoise fecal samples, we analyzed FASTq files from the Sequence Read Archive (Leinonen et al., 2011) database from tortoise fecal samples in EZBioCloud’s microbiome taxonomic profiling pipeline, using the PKSSU4.0 taxonomic identity database (Yoon et al., 2017).

### FASTq Files

All fecal sample FASTq files acquired from this study were uploaded to the SRA Database under the BioProject accession ID PRJNA967813.

## Results

### Analysis of Sequencing Results

After bioinformatics processing with the ZymoBiomics® targeted sequencing service for microbial identification and analysis, across 26 samples, a total of 9,305,862 raw sequences were read (Supplementary Table 1). After size filtration, 3,922,265 quality reads were obtained, totaling 25,912 final unique sequences. Final unique sequences ranged from 685 to 1,381 reads (Supplementary Table 1). All samples were rarefied to 20,000 reads. Rarefaction curves plateaued at a satisfactory sequencing depth across Shannon’s index (Figure 1) and other metrics/indices (Simpson’s, Simpson’s Reciprocal, Simpson’s e, PD whole tree, Chao1, Fisher alpha, observed species; Supplementary Tables 2A-G), suggesting microbial diversity among samples is representative. This suggests that a plausible approximation of within-sample diversity was achieved.

**Figure 1.**
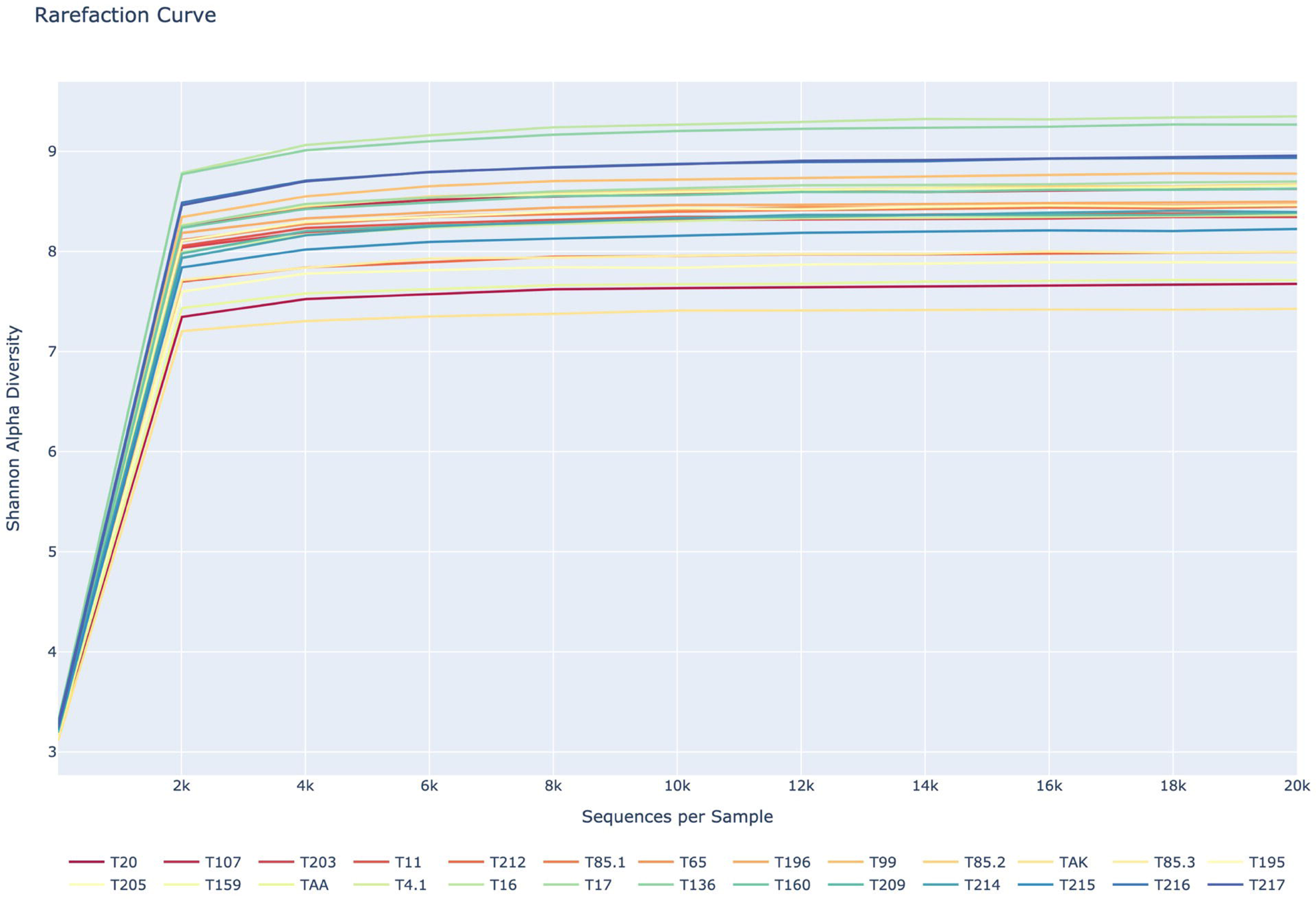
Rarefaction curves for each sample. The Shannon diversity scores for each sample plateau well before 20,000 sequences.

### Microbial Composition

The ZymoBiomics® taxonomic classifier identified 20 phyla (70.44% Firmicutes, 8.41% Bacteroidetes, 4.91% Euryarchaeota, and 7.57% other) (Figure 2), 37 classes (69.54% Clostridia, 8.41% Bacteroidia, 3.09 Methanobacteria, 1.91% Synergistia, and 1.49% Methanomicrobia), 52 orders (69.53% Clostridiales, 8.41% Bacteroidiales, and 3.09% Methanobacteriales), 78 families (27.49% Lachnospiraceae, 13.06% Ruminococcaceae, 11.68% Clostridiaceae, 7.73% Christensenellaceae, 6.79% of an unnamed Clostridiales family, 4.95% Rikenellaceae, 3.08% Methanobacteriaceae), 181 genera (18.29% of an unnamed Lacnhospiraceae genus, 11.68% *Clostridium*, 9.22% of an unnamed Ruminococcaceae genus, 7.73% of an unnamed Christensenellaceae genus, 4.68% of an unnamed Rikenellaceae genus, 4.23% *Cellulosilyticum,* and 4.05% of an unnamed Clostridiales family and genus), and 1509 “species” or operational taxonomic units (OTUs) (7.65% of an unnamed Lachnospiraceae genus and species sp32886, 5.09% *Clostridium butyricum,* and 4.69% *Clostridium* sp30783). See Supplementary Table 3B for all OTU/ASV assignments. OTUs are taxonomic assignments inferred from a reference library, in this case, given a ≥ 97% similarity to a taxon based on the sequenced amplicon sequence variants (ASV). The ASV that shows the highest similarity to a reference ASV with a known taxonomic assignment in the database is assigned that taxonomy.

**Figure 2.**
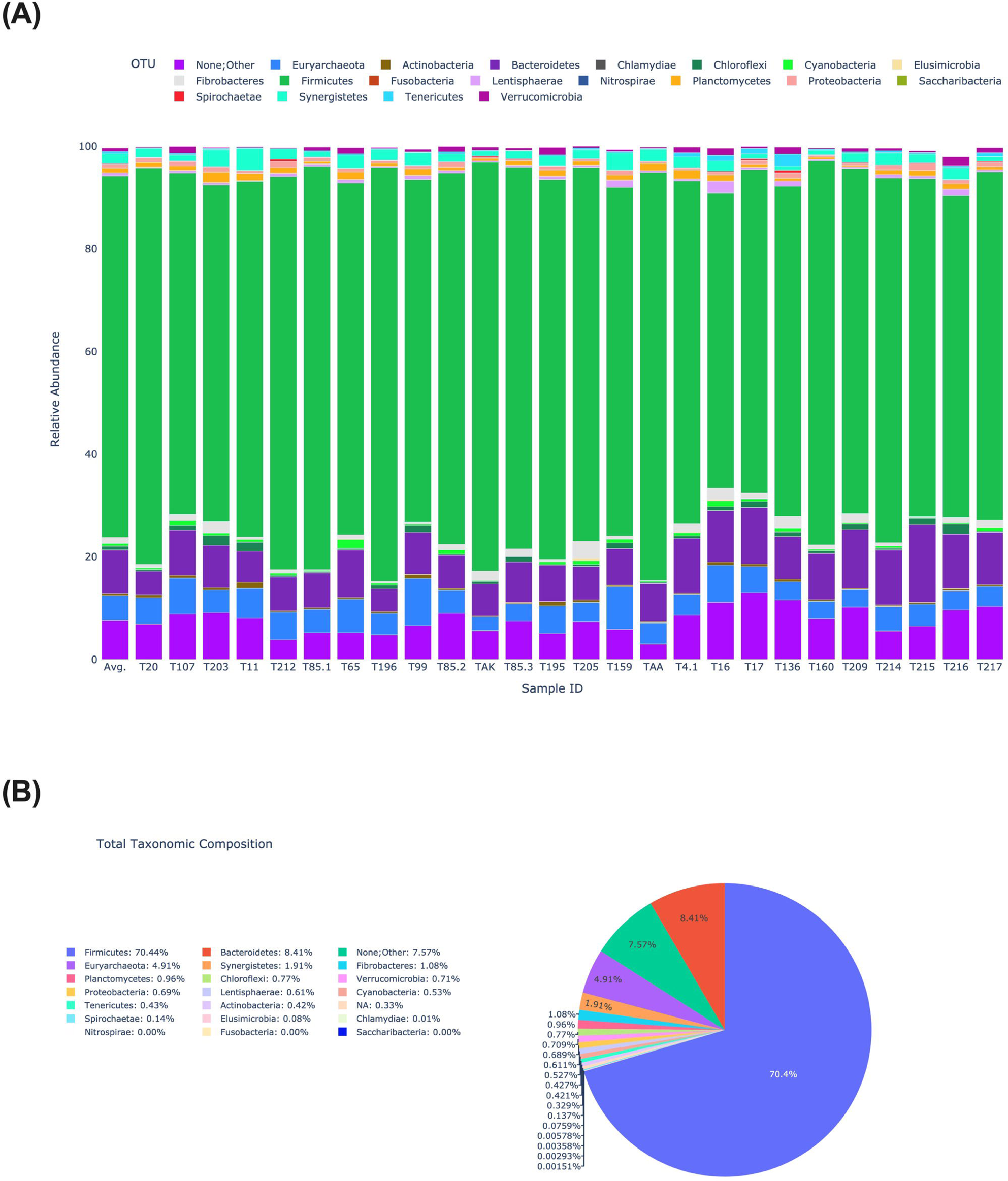
Plots representing relative abundance of phyla within each sample, average across samples, and core microbes across samples. **(A)** Bar plot representing average relative abundance phyla across samples, and within individual samples. Amplicon sequence variants (ASV) not assigned were categorized as “None;Other”. **(B)** Pie graph of community microbiome.

### Alpha Diversity

Comprehensive data for alpha diversity is in the supplementary materials (Supplementary Table 2). Overall, alpha diversity was high for our samples across indices. Observed OTUs [mean = 980.2, range = ± 650]; Shannon index [mean = 8.289, range = ± 1.514]; and Simpson index [mean = 0.991, range = ± 0.0193] (Supplementary Figure 1). No significant gender differences in alpha diversity were found across indices (Shannon, Observed OTUs, Chao1), with Mann-Whitney U tests showing *p* > 0.05 for all (Supplementary Figure 2).

### PCoA and Biplot

PC1 and PC2 of the PCoA explain 19.87% and 10.01% of the Bray-Curtis dissimilarity, respectively, across all samples (Figure 3A). PC1 and PC2 of the biplot explain 20.59% and 9.163% of the Bray-Curtis dissimilarity, respectively, across male and female samples, respectively (Figure 3A). The top 5 ASV loadings that explain the most dissimilarity are represented by the red arrows. The taxonomic assignments of the top 5 explanatory ASVs are *Clostridium butyricum*, Lachnospiraceae sp32886, Lachnospiraceae sp32886, Lachnospiraceae sp33372, and sequence/ASV 4.

**Figure 3.**
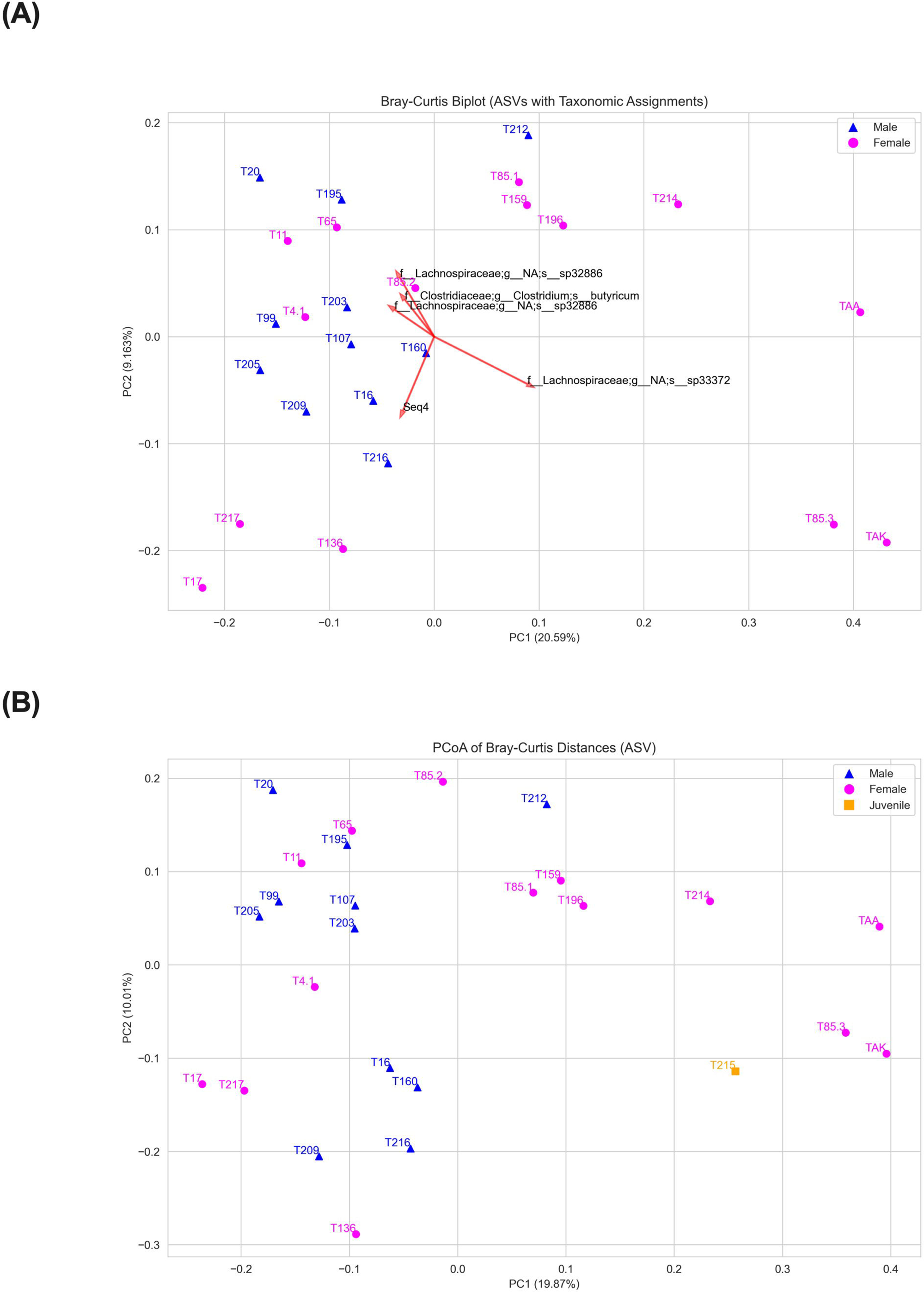
**(A)** Bray-Curtis biplot/principal coordinates analysis (PCoA) plot based on ASVs with taxonomic assignments. Each sample (only samples from sexed animals are shown) has a unique profile of ASV presence and abundance. This plot distinguishes dimensions that explain the dissimilarity between each profile. The PC axes show the chief dimensions that explain the dissimilarity between each sample. The red arrows represent the top five loadings (in this case ASVs) that linearly correlate with each presented PC axes. The taxonomy of each AVS is shown starting with family. The ASV, seq4, is 90.16% similar according to a BLASTn query, to “Uncultured Spirochaetaceae bacterium clone CH2 PE 73”. **(B)** PCoA plot with juvenile included.

### Beta Diversity

The dispersions for each iteration of the PERMDISP groups were not statistically different from one another, *p* > 0.05 (Supplementary Figure 3). The variances in beta diversity between males in females were only statistically significant different from one another when all temporally independent samples from the same tortoise T85 (i.e., T85.1, 2, and 3; Figure 4) were included, *p* < 0.05, PERMANOVA.

**Figure 4.**
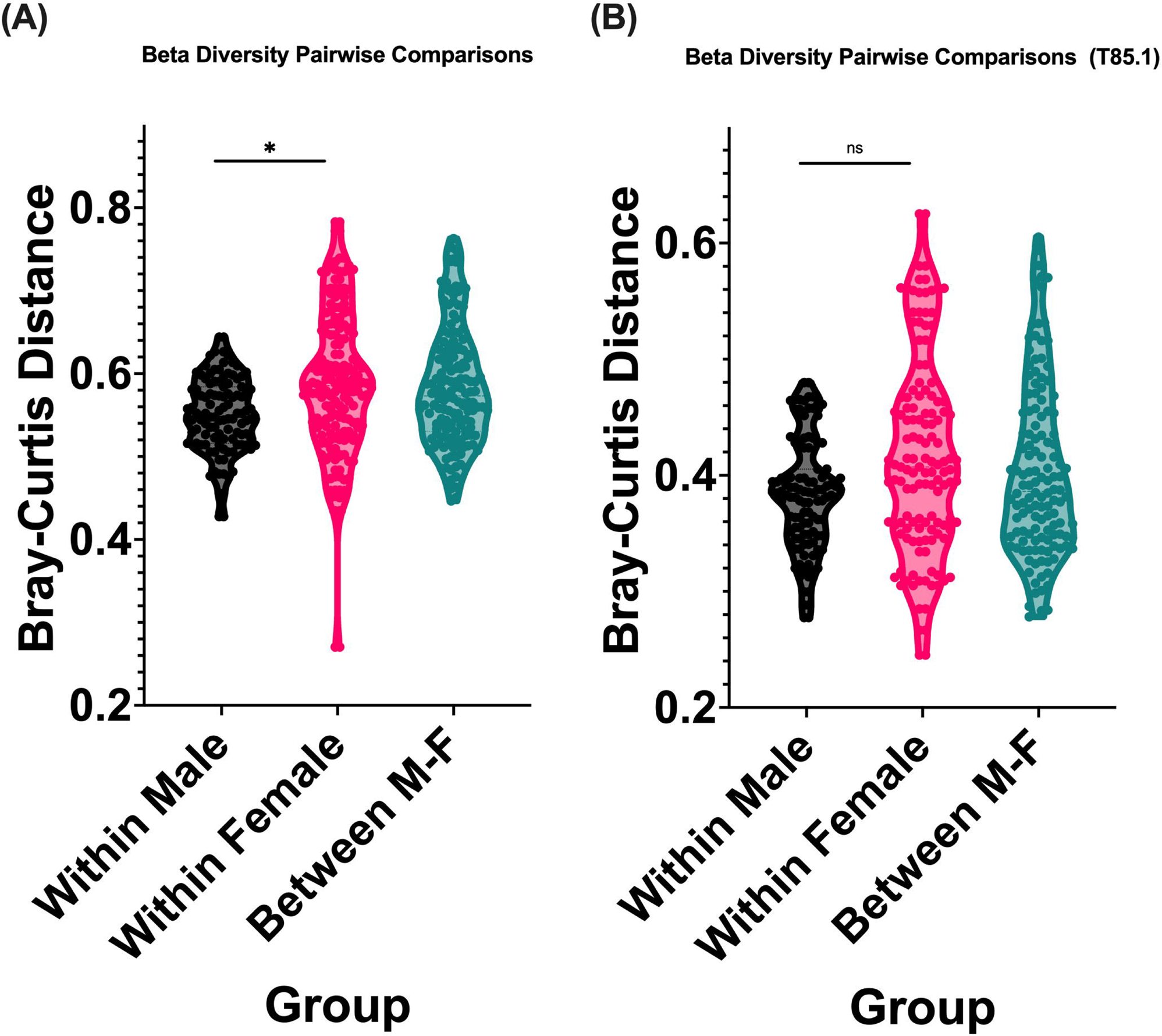
Bray-Curtis number of permutations = 999; * = *p* < 0.05, ns = not significant *p* > 0.05. **(A)** Male and female samples: PERMDISP *p* = 0.098, PERMANOVA *p* = 0.038, *R*^2^ = 6.60%. **(B)** Males and females. Females defined with just one temporally independent sample, T85.1: PERMDISP *p* = 0.274, PERMANOVA *p* = 0.217.

### Differential Abundance: LEfSe Analysis

The LEfSe (Linear discriminant analysis Effect Size) tool, v.1.1.2, was used to identify features that are statistically different between male and female samples (Figure 5). The analysis was conducted with a logarithmic LDA score threshold of 2.5 for discriminative features. The LEfSe analysis identified a total of 63 significantly discriminative features before the internal Wilcoxon test. Out of these, 15 features had an absolute LDA score > 2.5, indicating the strongest discriminations between the classes.

**Figure 5.**
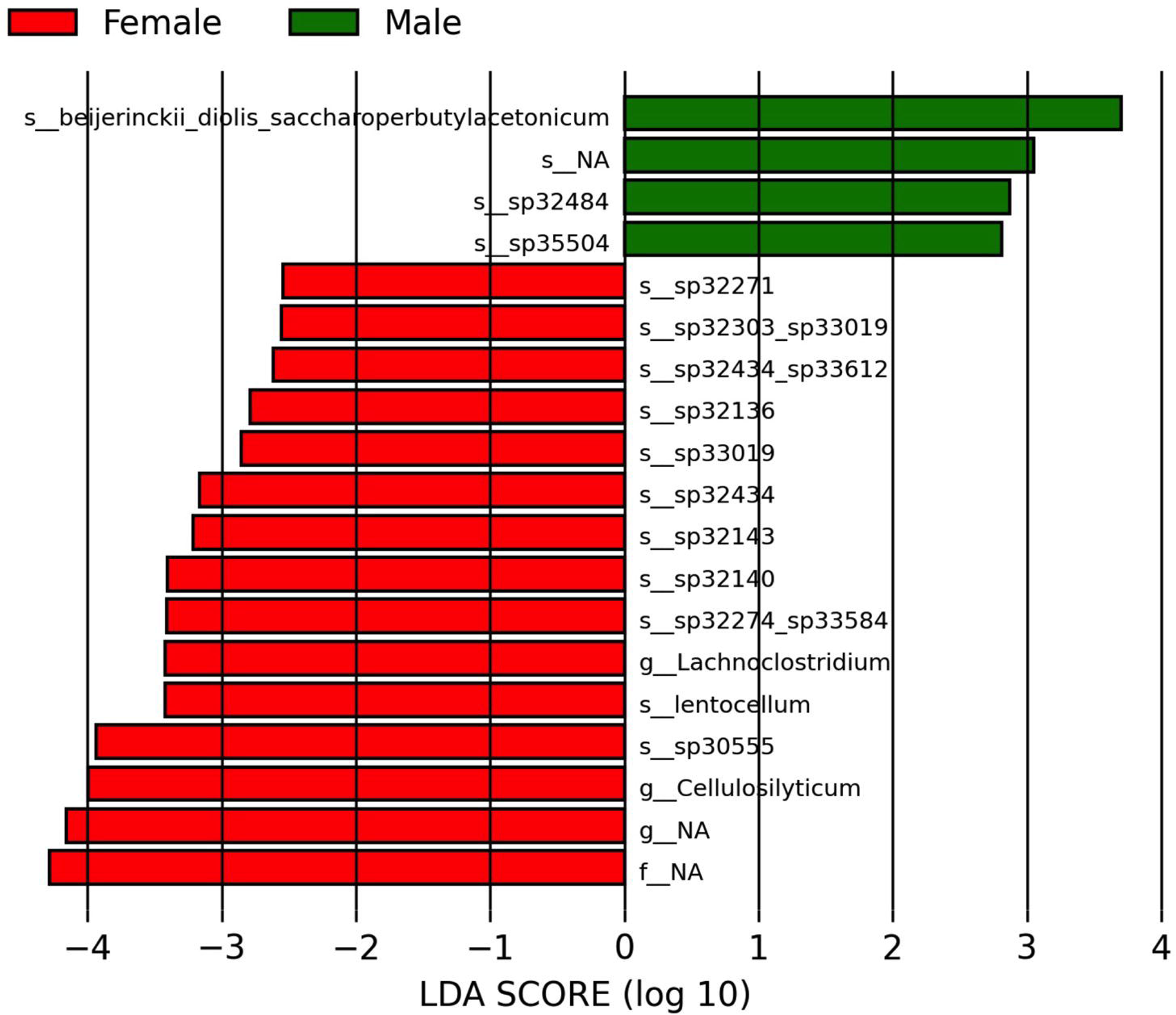
Linear discriminant analysis Effect Size (LEfSe). The waterfall plot shows differentially abundant taxa between sexes. LDA score threshold: 2.5 for discriminative features. Female bars point to the left and male bars point to the right. Taxonomic names have been shortened for ease of viewing. See Supplementary Table 4 for full names.

### Heatmap

Heatmaps show the differential abundances of top 1% of OTUs and ASVs between male and female samples (Figure 6). They corroborate biplot and LEfSe findings (Figures 3A and 6) and provide a more granular visualization of differential abundance and explanatory features across our samples.

**Figure 6.**
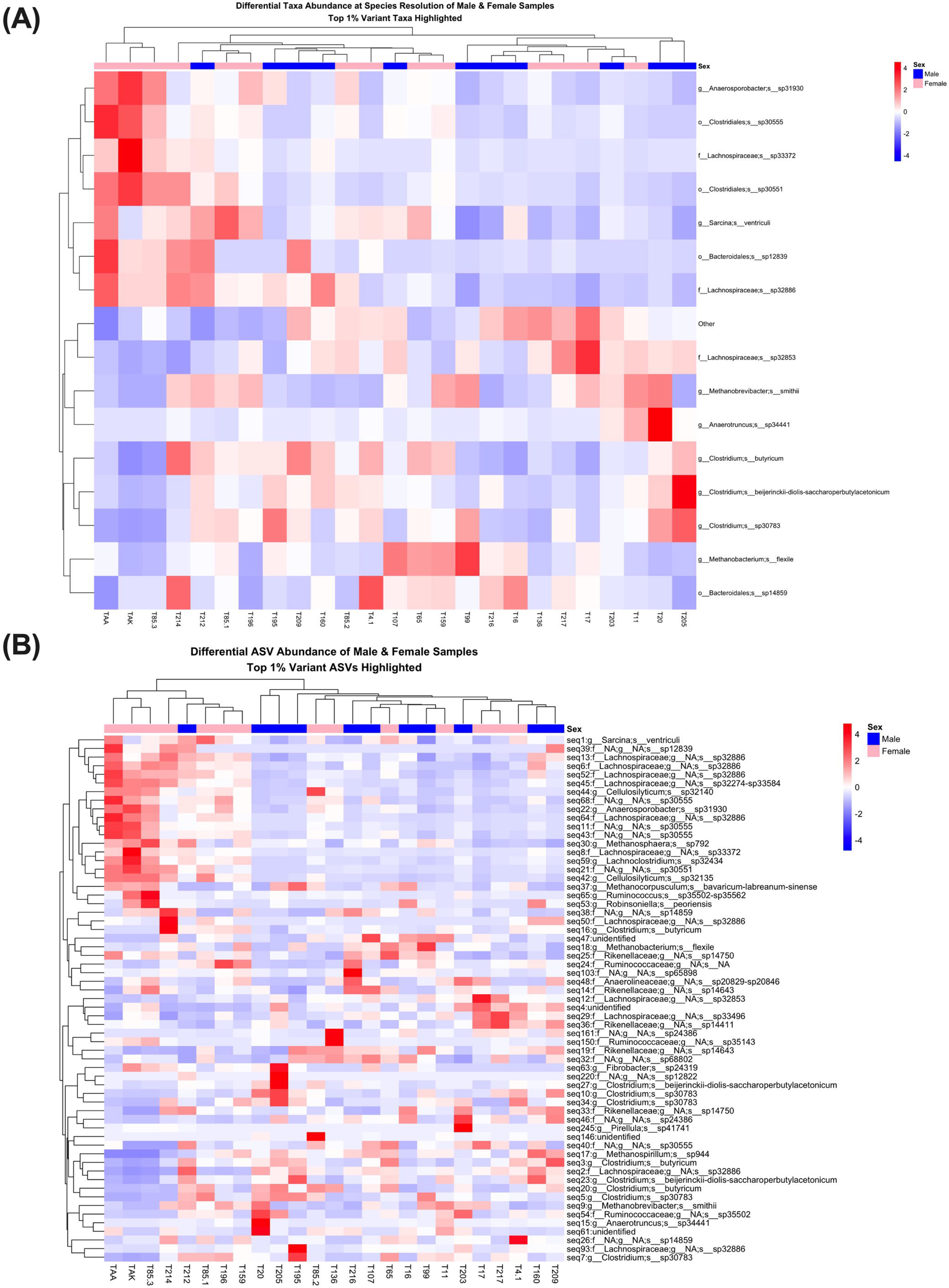
Heatmaps with clustering. Male and female samples are clustered together based on their relative feature abundances. The top 1% of differentially abundant features are presented. **(A)** Top 1% of OTUs at species resolution. **(B)** Top 1% ASVs at species resolution with taxonomic assignments.

**Table 1.**
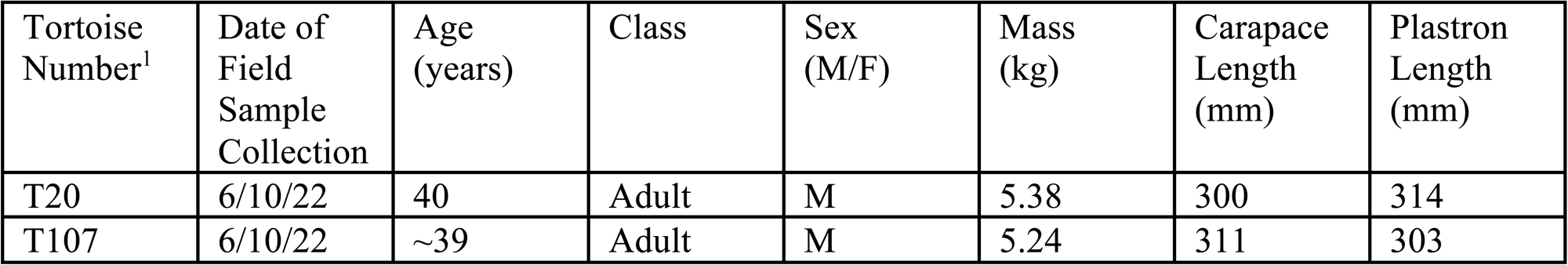
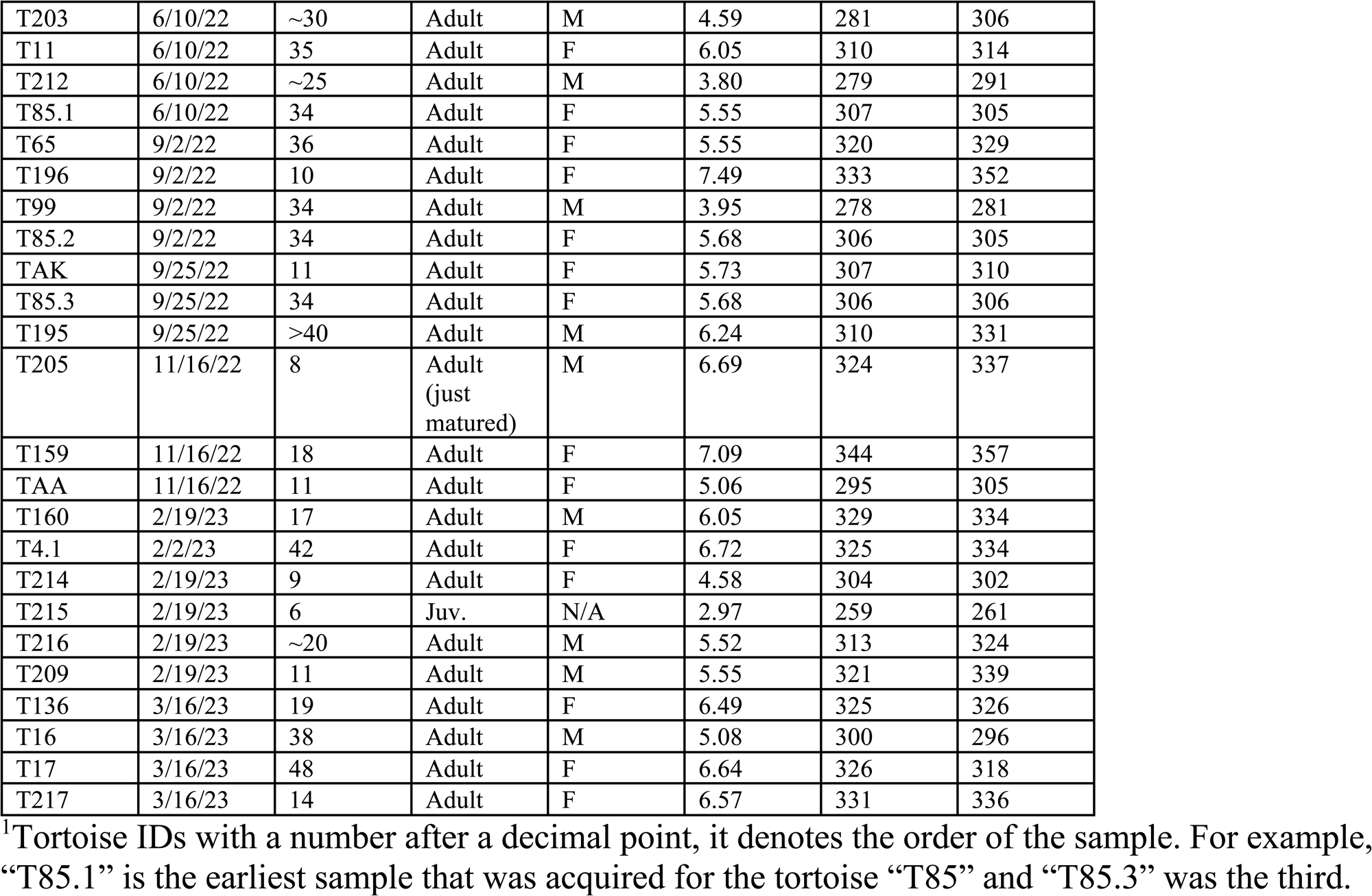
Gopher tortoise characteristics and sample metadata.

| Tortoise Number<sup>1</sup> | Date of Field Sample Collection | Age (years) | Class | Sex (M/F) | Mass (kg) | Carapace Length (mm) | Plastron Length (mm) |
| --- | --- | --- | --- | --- | --- | --- | --- |
| T20 | 6/10/22 | 40 | Adult | M | 5.38 | 300 | 314 |
| T107 | 6/10/22 | ~39 | Adult | M | 5.24 | 311 | 303 |
| T203 | 6/10/22 | ~30 | Adult | M | 4.59 | 281 | 306 |
| T11 | 6/10/22 | 35 | Adult | F | 6.05 | 310 | 314 |
| T212 | 6/10/22 | ~25 | Adult | M | 3.80 | 279 | 291 |
| T85.1 | 6/10/22 | 34 | Adult | F | 5.55 | 307 | 305 |
| T65 | 9/2/22 | 36 | Adult | F | 5.55 | 320 | 329 |
| T196 | 9/2/22 | 10 | Adult | F | 7.49 | 333 | 352 |
| T99 | 9/2/22 | 34 | Adult | M | 3.95 | 278 | 281 |
| T85.2 | 9/2/22 | 34 | Adult | F | 5.68 | 306 | 305 |
| TAK | 9/25/22 | 11 | Adult | F | 5.73 | 307 | 310 |
| T85.3 | 9/25/22 | 34 | Adult | F | 5.68 | 306 | 306 |
| T195 | 9/25/22 | >40 | Adult | M | 6.24 | 310 | 331 |
| T205 | 11/16/22 | 8 | Adult<br>(just<br>matured) | M | 6.69 | 324 | 337 |
| T159 | 11/16/22 | 18 | Adult | F | 7.09 | 344 | 357 |
| TAA | 11/16/22 | 11 | Adult | F | 5.06 | 295 | 305 |
| T160 | 2/19/23 | 17 | Adult | M | 6.05 | 329 | 334 |
| T4.1 | 2/2/23 | 42 | Adult | F | 6.72 | 325 | 334 |
| T214 | 2/19/23 | 9 | Adult | F | 4.58 | 304 | 302 |
| T215 | 2/19/23 | 6 | Juv. | N/A | 2.97 | 259 | 261 |
| T216 | 2/19/23 | ~20 | Adult | M | 5.52 | 313 | 324 |
| T209 | 2/19/23 | 11 | Adult | M | 5.55 | 321 | 339 |
| T136 | 3/16/23 | 19 | Adult | F | 6.49 | 325 | 326 |
| T16 | 3/16/23 | 38 | Adult | M | 5.08 | 300 | 296 |
| T17 | 3/16/23 | 48 | Adult | F | 6.64 | 326 | 318 |
| T217 | 3/16/23 | 14 | Adult | F | 6.57 | 331 | 336 |

<sup>1</sup>Tortoise IDs with a number after a decimal point, it denotes the order of the sample. For example, “T85.1” is the earliest sample that was acquired for the tortoise “T85” and “T85.3” was the third.

### Potential Pathogens and Probiotics

Prospective gopher tortoise probiotics we detected in our samples included Lachnospiraceae, *Clostridium butyricum*, *Faecalibacterium prausnitzii*, and *Ruminococcus bromii* (Ariyoshi et al., 2022; Huang et al., 2019; Lopez-Siles et al., 2017; Ze et al., 2012). Potential pathogens detected in our samples included Mycoplasmaceae sp., *Helicobacter* sp.; potential pathogens not detected in our samples included Gopher Tortoise *Helicobacter, Anaplasma* spp., and *Mycoplasma* spp. (Gilbert et al., 2014, 2019; Raskin et al., 2020; Desiderio et al., 2021; Page-Karjian et al., 2021)(Table 2).

**Table 2.** Potential pathogens and probiotics. This table depicts potential probiotics present in our samples, and potential pathogens detected in our samples. (+) = detected (-) = not detected.

| Potential Probiotics | Potential pathogens |
| --- | --- |
| Lachnospiraceae (+) | Mycoplasmaceae (+) |
| *Clostridium butyricum* (+) | *Helicobacter sp55646* (+) |
| *Faecalibacterium prausnitzii* (+) | Gopher Tortoise *Helicobacter* (-) |
| *Ruminococcus bromii* (+) | *Anaplasma spp.* (-) |
|  | *Mycoplasma testudineum* (-) |
|  | *Mycoplasma agassizii* (-) |

### Post Hoc Taxa Investigation and Analyses

The *H.* sp55646 (seq1479) ASV present within (*n* = 6) tortoises (*n* = 6 fecal samples) (Supplementary Table 3A) was 97% similar to GT *Helicobacter* (accession ID: MT758473.1); and *H.* sp55646 had a 100% pairwise similarity to *Helicobacter* sp. 12S02256-12 (Accession ID: KJ081208.1) (Supplementary Data 1).

The OTU, Mycoplasmataceae sp68290, was present in two fecal samples from two tortoises. The associated ASV, seq2192, according to a BLAST query, had a 90-95.75% sequence similarity to *Mycoplasma* and uncultured bacterial species (Supplementary Data 2).

*Mycoplasma* spp. rRNA genes can be found in tortoise fecal samples. *Mycoplasma* spp. were present in five tortoises (SRA run IDs: ERR4994823, SRR5190049, ERR3988747_1, ERR4995630, ERR3672735; the first four runs are from Seychelles giant tortoises (*Aldabrachelys gigantea hololissa*), and the last one is from a Burmese star tortoise (*Geochelone platynotan*) (Supplementary Data 3).

## Discussion

The fecal microbiomes of the Abacoa Greenway Range VIa gopher tortoise population were mainly comprised of the phyla Firmicutes (70.4%), Bacteroidetes (8.41%), and Euryarchaeota (4.91%). The Clostridia class comprised 69.54% of the gut microbiome. Notable families included 27.49% Lachnospiraceae, 13.06% Ruminococcaceae, and 11.68% Clostridiaceae (many taxa within these families are not named by the taxonomic database implemented, possibly due to the lack of studies on the reptile gut microbiome). 6,025 ASVs and 1,509 OTUs were recovered, signifying a high level of alpha diversity.

Overall, these findings agree with previous investigations in the fecal microbiomes of herbivorous reptiles (Siddiqui et al., 2022; Hoffbeck et al., 2023) and mammals (Zhu et al., 2021; Wu et al., 2022). Firmicutes and Bacteroidetes are phyla known to play a role in the metabolism of dietary starches, e.g., cellulose and hemicellulose, which make up the cell walls of plant cells, into SCFAs. Butyrate, a remarkable SCFA, plays a role in immune function (Siddiqui et al., 2022), gut physiology (Koh et al., 2016; Carretta et al., 2021; van der Hee and Wells, 2021), epithelial integrity (Braniste et al., 2014; Parada Venegas et al., 2019), and vagal nerve signaling (Cawthon and de La Serre, 2018). Moreover, gopher tortoises are known as hind-gut-fermenters, for which these starch fermenting bacteria are responsible. Hence, uncovering the significant roles these unnamed taxa likely play within herbivore gut microbiome is warranted.

In terms of microbial composition, findings from previous gopher tortoise gut microbiome studies were remarkably different compared to ours. Gaillard (2014) reported an overall average abundance of 59.7% Firmicutes, 15.9% Bacteroidetes, 15.4% Proteobacteria, and 1.9% Planctomycetes (Gaillard did not screen for Euryarchaeota; this may confound cross-study comparisons), while Yuan et al. (2015) uncovered 36.5% Bacteroidetes, 36% Firmicutes, 3.13% Verrumicrobia, 1.17% of Euryarchaeota, 1.16% Proteobacteria, and 0.6% of Planctomycetes. We found 70.4% Firmicutes, 8.41% Bacteroidetes, 4.91% Euryarchaeota 0.96% Planctomycetes, 0.71% Verrumicrobia, and 0.69% Proteobacteria. It is difficult to explain, however, differences in microbial composition in tortoises across these studies.

Relatedly, wild and captive populations of *Gopherus flavomarginatus*, the most phylogenetically related species to the gopher tortoise (Thomson et al., 2021), had microbiomes made up of an average of 86% Firmicutes (Garcia-De La Peña et al., 2019). However, interestingly, 0% Bacteroidetes were detected (Garcia-De La Peña et al., 2019). Further, populations of *Gopherus agassizii* had a microbiome composition dominated by Firmicutes and Bacteroidetes, respectively (Drake et al., 2017). This suggests the ubiquity of Firmicutes within *Gopherus* spp. Moreover, it is curious why *G. flavomarginatus* lacked Bacteroidetes given its commonality in turtles (Yuan et al., 2015; Abdelrhman et al., 2016; Modica, 2016; Ahasan et al., 2017, 2018, 2020; Campos et al., 2018; Arizza et al., 2019; Garcia-De La Peña et al., 2019; Bloodgood et al., 2020; Fong et al., 2020, 2024; Fugate et al., 2020; McDermid et al., 2020; McKnight et al., 2020; Qu et al., 2020; Sandri et al., 2020; Scheelings et al., 2020b, 2020a; Filek et al., 2021; Hong et al., 2021; Beale et al., 2022; Chen et al., 2022; Díaz-Abad et al., 2022; Giakoumas, in press; Khan et al., 2024; Niu et al., 2024) and herbivores (Zhu et al., 2021; Zoelzer et al., 2021; de Jonge et al., 2022; Wu et al., 2022; Niu et al., 2024).

### Proposed Explanations for Microbial Differences Across *G. polyphemus* Populations

Yuan et al. (2015) reason that the significant (*prima facie*) difference in relative abundance (36.5%) of the Bacteroidetes phylum from their population in contrast to the northern gopher tortoise populations (15.9%), as investigated by Gaillard (2014), may have to do with the fact that their study populations are located within in different climates; they also mention the differences in Firmicutes and Bacteroidetes ratios in populations of herbivorous reptiles from more tropical regions. Considered together, they speculate that differences in Firmicutes and Bacteroidetes ratios may be due to differences in climate. Nevertheless, our population is located in a warmer, more tropical climate than the Archbold population, and has ∼70% Firmicutes and ∼15% Bacteroidetes, hence, confounding Yuan et al.’s hypothesis. In this respect, the Archbold population is unique compared to other gopher tortoise populations.

In general, for reptiles, Hoffbeck et al. (2023) report methodology as being the most influential, accounting for almost a quarter of variation of a host species’ microbiome composition. Methodological variables included were host extraction area, 16S rRNA hypervariable region, study, and DNA extraction method. DNA extraction method accounted for the most variation. Variables such as host diet, order, conservation status, and captivity status explain less than 1% of the variance in Bray-Curtis microbial dissimilarity. Across each *G. polyphemus* study, a different DNA extraction method was used; thus, DNA extraction method may be the salient variable explaining microbial composition variance across each *G. polyphemus* study.

### Males Verses Females Composition

Differential abundance analysis reveals there are no significant differences between male and female gut microbiome composition (Figure 4). However, differential abundance analysis reveals there are differentially abundant taxa among male and female samples, with a modest effect size. However, little research has been conducted on these biomarkers.

Counterintuitively, male taxonomic composition profiles were less dispersed than females (Figure 4). This is the case even though males tend to travel more often, longer distances, and have a larger home range than females (McRae et al., 1981b; Diemer, 1992). Conspecific overcrowding, geographic confinement (Sano, 2014; Jon Moore, personal communication, 2023), homogeneity of forage, and geographically conservative mating strategies may explain these counterintuitive results. TAA, TAK, and T85.3, all samples from females, cluster together across the two largest Bray-Curtis PCoA axes, and probably account for why there is a significant difference between male and female samples, but not males and females. One ASV, unstudied but classified as an unnamed species, Lachnospiraceae sp33372, may drive these differences, Figure 3A. One possible explanation for why these samples are distinguished from the rest may be related to a female gestation response: When female tortoises are producing the shells for their eggs, they often supplement their diet with calcium from stones, fossil shells, or small mammal bones from feces of coyotes, bobcats, or foxes (Moore and Dornburg, 2014).

### Potential Pathogens and Probiotics

#### Potential Pathogens

Gopher tortoises are commonly afflicted with upper-respiratory tract diseases (URTD) (Fremont, 2017; Raskin et al., 2020; Page-Karjian et al., 2021) and endoparasites (Huffman et al., 2018; Cooney et al., 2019), which may hinder conservation efforts. *Mycoplasma* (Page-Karjian et al., 2021) and *Anaplasma* (Raskin et al., 2020; Page-Karjian et al., 2021) genera are common culprits for URTD infections; and a novel *Helicobacter* species, provisionally dubbed “gopher tortoise *Helicobacter*” (GT *Helicobacter*), is a suspected culprit for three, and potentially more, gopher tortoise infections (Desiderio et al., 2021). Thus, we screened for these potential pathogens in our sample population (Table 2).

*Helicobacter* spp. are commonly pathogenic species with the ability to afflict a broad phylogeny of eumetazoans (Gilbert et al., 2014, 2019). However, in reptiles, the clinical relevance of *Helicobacter* species to tortoise health is understudied. Most notably, the novel *Helicobacter* strain, “gopher tortoise *Helicobacte*r (GT *Helicobacter*)”, uncovered by Desiderio et al. (2021) has been shown to be associated with gopher tortoise mortality. In ill tortoises, Desiderio et al. (2021) took nasal swabs and sequenced the whole 16S rRNA gene (accession ID: MT758473.1) and two other highly conserved genes in order to identify the novel strain. Relatedly, in our population, we found one *Helicobacter* OTU present within (*n =* 6) tortoises (*n* = 6 fecal samples): *Helicobacter* sp55646; one V3-V4 16S sequence was associated with this *Helicobacter* OTU. When analyzed with BLAST (Zhang et al., 2000; Morgulis et al., 2008) it had a 100% alignment match with *Helicobacter* sp. 12S02256-12 (Accession ID: KJ081208.1). This species was found in the Warren’s girdled lizard (*Cordylus warren*) (Gilbert et al., 2014). This lizard is native to southern Africa. Relatedly, an invasive species of lizard, African redhead agama (*Agama picticauda*), in south Florida (Gray, 2020; Mitchell et al., 2021) has been observed at Florida Atlantic University’s Harriet L. Wilkes Honors College (personal observation, Giakoumas, 2022-2024) which is located near Range VIa of the Abacoa Greenway. It is possible that this species of *Helicobacter* was transmitted from an African lizard to the gopher tortoises of this study via coprophagy.

Additionally, *Mycoplasma* spp. and *Anaplasma* spp. cause URTD (Fremont, 2017; Raskin et al., 2020; Page-Karjian et al., 2021) in gopher tortoises; more specifically, *M. agassizii* have been found to afflict *Gopherus* spp. (Fremont, 2017; Page-Karjian et al., 2021). *Anaplasma* spp., were not detected in the feces of our samples. Interestingly, one OTU, sp68290, seq2192, which was associated with the family Mycoplasmataceae (Supplementary Table 3), was present in two fecal samples from two tortoises. The associated ASV, seq2192, has a 90-95.75% sequence similarity to *Mycoplasma* and uncultured bacterial species, according to a BLAST query (Supplementary Data 2). Although this ASV is phylogenetically distinct from known pathogenic *Mycoplasma* spp., it is categorized within the class, Mollicutes, which are difficult to treat with antibiotics (Chernova et al., 2021). Therefore, further investigation of the potential pathogenicity of this ASV is still warranted.

Taken together, the lack of detection of known or suspected pathogenic *Mycoplasma*, *Anaplasma*, and GT *Helicobacter* provides a line of evidence against the hypothesis that these taxa are present in our population. Despite this, since these taxa more prominently infect the respiratory tract in gopher tortoises (Desiderio et al., 2021; Page-Karjian et al., 2021), they may be detected via nasal swab. Therefore, nasal probes should be employed to further test this hypothesis. Just because these genera are not known to play an etiological role in the gopher tortoise gastrointestinal tract, does not mean their presence in the gastrointestinal tract is irrelevant. *H. pylori*, for instance, can survive the gastrointestinal tract (Yang et al., 2013; Malfertheiner et al., 2023); in fact, there is evidence to suggest that GT *Helicobacter* can be present in gopher tortoise fecal samples and that *Helicobacter* can be found in gopher tortoise fecal samples within a week after defecation *in situ et in natura* (Giakoumas, unpublished data). So, it may be the case that gopher tortoises obtain GT *Helicobacter* via coprophagy. Additionally, the many invasive reptilian species in South Florida (Searcy et al., 2023) may jeopardize gopher tortoise health if their feces are consumed.

#### Potential Probiotics

Among the notable potential probiotic microbes we detected were those within the family Lachnospiraceae, and in the genera *Clostridium*, *Ruminococcus*, and *Faecalibacterium*. These taxa are all subcategorized in the phylum Firmicutes. As expected, since most known microbial species that have been studied have been studied because they have some relationship to humans, many OTUs detected within our gopher tortoise population are unnamed or understudied. Still, we can make inferences from broad generalizations about their functions in relation to the gopher tortoise for each taxon.

The Lachnospiraceae family is mainly associated with studied pathologies that pertain to humans (Vacca et al., 2020); these pathologies are of little relevance in gopher tortoises, however. Nevertheless, there are probiotic taxa from the Lachnospiraceae family that are associated with SCFA production, as many Firmicutes do, such as butyrate, propionate, and acetate (Zaplana et al., 2024). This is not surprising, as gopher tortoises consume mainly vegetation, fueling the metabolism of SCFA from undigestible starches via bacterial fermentation. It is even emerging as a potential biotherapeutic in humans (Zaplana et al., 2024). Notably, the OTU “sp32886” detected within our samples of the Lachnospiraceae family had the largest average relative abundance at 7.65%. This far exceeds the relative abundances of almost all the other species level OTUs. It is worth, therefore, further investigation and characterization.

*Ruminococcous bromii* was detected in 17 samples with a total average relative abundance of 0.03%. This is remarkable as *R. bromii* is a keystone species in SCFA production in the human gut microbiome (Ze et al., 2012; Crost et al., 2018). This species initiates the breakdown of indigestible starches, particularly alpha-(1-4)-linked oligosaccharides, supports gut probiotic community members, and increases the amount of butyrate production (Ze et al., 2012). However, *R. bromii* was only present in small amounts (range = 0.138%) across all of our samples. Given that gopher tortoises are hindgut fermenters, this should prompt further investigation into which, if any, butyrate up-regulating keystone taxa specifically inhabit the gopher tortoise.

Perhaps the most notable OTU that was detected within our population was *Clostridium butyricum*. The average relative abundance of *C. butyricum* was 5.09% and was present across all samples (minimum 1.63% and maximum 9.93%). *C. butyricum*, besides being responsible for SCFA production, seems to have the ability to combat a few pathogenic bacteria (Ariyoshi et al., 2022). The *C. butyricum* strain, MIYAIRI 588 (CBM 588), for example, has been shown to be linked to the elimination of *H. pylori* in the gastrointestinal tract (Takahashi et al., 2000). This relationship, however, may be explained by the activation of a systemic immune response by CBM 588 and other non-respiratory specific mechanisms (Ariyoshi et al., 2022). Thus, *C. butyricum* may not be directly applicable to GT *Helicobacter* treatment, unless the pathogenesis of GT *Helicobacter* involves gastrointestinal colonization, which is plausible as a *Helicobacter* species appears in turtle fecal samples (Supplementary Table 3A; Giakoumas, unpublished data).

Many more bacteria within our Abacoa population likely have unique symbiotic functions beyond mere SCFA production via fermentation of plant fibers; however, most probiotic research has been conducted on mammals. Additionally, it is possible that certain potential pathogens may have gone undetected, and thus, a discussion about species level activity in reptiles is significantly limited. Further, 16S taxonomic classification is not granular enough to appreciably predict strain or species level functions; whole-genome shotgun sequencing and analysis, and metabolomic analysis would be helpful here. Nevertheless, this study provides a foundation for better understanding the host-microbiome linkages in this threatened keystone reptile.

## Data Availability Statement

The datasets presented in this study can be found in online repositories. The names of the repository/repositories and accession number(s) can be found below: All fecal sample FASTq files acquired from this study were uploaded to the SRA Database under the BioProject accession ID PRJNA967813, reviewer link: <u>to non-reviewers, FASTq files available upon</u> <u>request.</u>

## Ethics Statement

All capture, handling, and sampling procedures were approved by the following: Florida Atlantic University’s Institutional Animal Care and Use Committee (IACUC) approvals (A-19-41; A-22-47; A(T)22-06) and Florida Fish and Wildlife Conservation Permit (LSSC-14-00066B & C).

## Author Contributions

DG, TM, ES conceived of and designed the study. DG, ES, and KB acquired grants and funding. DG carried out the research. DG and JM collected fecal samples. JM physically assessed tortoises. DG extracted DNA from fecal samples. Zymo Research Corporation performed next-generation sequencing and bioinformatic processing and analysis on samples. DG performed bioinformatics processing, analysis, and visualization. TM, ES, KB, and JM edited and reviewed the manuscript. DG wrote most of the manuscript. JM contributed a few paragraphs with expertise on the gopher tortoises and the Abacoa Greenway. ES, KB, and TM supervised, managed, and performed administrative duties. KB and ES provided laboratory materials and space. All authors contributed to the article and approved the submitted version.

## Acknowledgments

We thank the Dean, Justin Perry, of the Florida Atlantic University Harriet L. Wilkes Honors College, and the Florida Atlantic University Office of Undergraduate Research and Inquiry for funding our research. We acknowledge that this research article builds upon Giakoumas’ honors thesis research at the Harriet L. Wilkes Honors College, whereby 10 more fecal samples were acquired, sequenced, processed and analyzed, and further analyses were conducted; this thesis is in press as of the current date of submission. We would like to thank Morgan Slevin for meeting early on for advice on conducting a microbiome study. Additionally, we thank Zymo Research for providing Zymo Research’s DNA extraction kits and microBIOMICS sequencing and analysis service. We thank Dr. Luisa Galgani for her recommendations for contacts at Zymo Research. We thank Dr. David Bradshaw, Dr. Alyssa Demko, and Dr. Ryan Bos for creating and/or providing us with coding materials and insight into microbiome analysis. We thank Ashley Boswell for assistance in DNA extraction. We thank Dr. Iris Segura-García for her supervision and counsel on microbiome research analysis. We acknowledge that ChatGPT-4 was used to generate template scripts for data visualization and analysis.

## Supplementary Material

The Supplementary Material for this article can be found online at: scripts used by the author in this study are available here: https://github.com/dgiakou/GT_Gut_Microbiome_Abacoa_2024.git

## Supplementary Material

### Supplementary Figures

**Supplementary Figure 1.** Alpha diversity indices per sample.

**Supplementary Figure 2.** Alpha diversity box plots for male and female. “+” represents mean. No significant difference was found between groups.

**Supplementary Figure 3.** Violin plots of Bray-Curtis distances with each temporally independent sample from T85. Number of permutations = 999; * = *p* < 0.05, ns = not significant *p* > 0.05. No significant difference between males and females regardless of each T85 sample, according to PERMANOVA tests. **(A)** T85.1: PERMDISP *p* = 0.274; PERMANOVA *p* = 0.217. **(B)** T85.2: PERMDISP *p* = 0.314; PERMANOVA *p* = 0.256. **(C)** T85.3: PERMDISP *p* = 0.120; PERMANOVA *p* = 0.163.

