## Supplementary_Data_1 for "State-Threatened Gopher Tortoise (*Gopherus polyphemus*) Gut Microbiome Analysis Reveals Health Insights into Southeastern Florida Population"

```
RID: 6CU7DWWS016
Job Title:seq1479
Program: BLASTN
Database: nt Nucleotide collection (nt)
Query #1: Uncultured Helicobacter sp. clone R16-645 16S ribosomal RNA gene, partial sequence Query ID: gb|MT758473.1 Length: 1391

Sequences producing significant alignments:
                                                                  Scientific      Common                     Max    Total Query   E   Per.   Acc.
Description                                                       Name            Name            Taxid      Score  Score cover Value Ident  Len        Accession
Uncultured Helicobacter sp. clone R16-645 16S ribosomal RNA...    uncultured H... NA              175537     2569   2569  100%  0.0   100.00 1391       MT758473.1
Helicobacter sp. 11S02629-2 16S ribosomal RNA gene, partial...    Helicobacter... NA              1476195    2200   2200  99%   0.0   95.33  1391       KJ081205.1
Helicobacter sp. strain MIT 16-1353 16S ribosomal RNA gene,...    Helicobacter... NA              218        2170   2170  99%   0.0   94.83  1504       MW147609.1
Helicobacter sp. 'pancake tortoise' 16S ribosomal RNA gene,...    Helicobacter... NA              611767     2169   2169  95%   0.0   96.23  1370       FJ667779.1
Helicobacter sp. MIT 13-11044 16S ribosomal RNA gene, partial...  Helicobacter... NA              1680556    2137   2137  100%  0.0   94.41  1470       KT159712.1
Helicobacter sp. MIT 13-11042 16S ribosomal RNA gene, partial...  Helicobacter... NA              1680555    2137   2137  100%  0.0   94.41  1470       KT159711.1
Helicobacter sp. 12S02256-12 16S ribosomal RNA gene, partial...   Helicobacter... NA              1476198    2135   2135  99%   0.0   94.40  1403       KJ081208.1
Helicobacter sp. strain MIT 16-1358 16S ribosomal RNA gene,...    Helicobacter... NA              218        2126   2126  98%   0.0   94.57  1444       MW147611.1
Helicobacter trogontum strain MIT 95-5368 16S ribosomal RNA...    Helicobacter... NA              50960      2126   2126  100%  0.0   94.28  1470       AY686609.1
Helicobacter trogontum strain LRB 8581 16S ribosomal RNA,...      Helicobacter... NA              50960      2126   2126  100%  0.0   94.28  1422       NR_026029.1
Helicobacter jaachi strain MIT 09-6949 16S ribosomal RNA,...      Helicobacter... NA              1677920    2124   2124  100%  0.0   94.21  1459       NR_149209.1
Helicobacter sp. 'Flexispira taxon 5' strain ATCC 43966 16S...    Helicobacter... NA              50960      2115   2115  100%  0.0   94.13  1446       M88137.2
Helicobacter sp. RTP4S 16S ribosomal RNA gene, partial sequence   Helicobacter... NA              930050     2113   2113  98%   0.0   94.48  1447       AY554133.1
Helicobacter sp. RTP3S 16S ribosomal RNA gene, partial sequence   Helicobacter... NA              930049     2113   2113  98%   0.0   94.48  1443       AY554132.1
Helicobacter sp. RTP2S 16S ribosomal RNA gene, partial sequence   Helicobacter... NA              930048     2113   2113  98%   0.0   94.48  1487       AY554131.1
Helicobacter sp. RTP5S 16S ribosomal RNA gene, partial sequence   Helicobacter... NA              930051     2109   2109  98%   0.0   94.42  1449       AY554134.1
Helicobacter sp. RTP1S 16S ribosomal RNA gene, partial sequence   Helicobacter... NA              930047     2104   2104  98%   0.0   94.35  1459       AY554130.1
Helicobacter sp. IMMIB HP-28/08 partial 16S rRNA gene, type...    Helicobacter... NA              524885     2098   2098  99%   0.0   94.16  1458       AM992061.1
Helicobacter sp. BTP2S 16S ribosomal RNA gene, partial sequence   Helicobacter... NA              281982     2098   2098  99%   0.0   93.92  1474       AY554136.1
Helicobacter canis isolate MGYG-HGUT-01463 genome assembly,...    Helicobacter... NA              29419      2097   4188  100%  0.0   93.84  1932823    LR698964.1
Helicobacter canis strain MIT 99-7633 16S ribosomal RNA gene,...  Helicobacter... NA              29419      2097   2097  100%  0.0   93.84  1473       KC878292.1
Helicobacter sp. strain 91-266-11 Fox 16S ribosomal RNA, parti... Helicobacter... NA              279279     2097   2097  100%  0.0   93.86  1449       M88152.2
Helicobacter canis 16S ribosomal RNA gene, partial sequence       Helicobacter... NA              29419      2097   2097  100%  0.0   93.84  1455       AF177475.1
Helicobacter sp. Dog-4 16S ribosomal RNA gene, partial sequence   Helicobacter... NA              279210     2091   2091  100%  0.0   93.78  1425       U51874.2
Helicobacter canis strain MIT 51402 16S ribosomal RNA gene,...    Helicobacter... NA              29419      2091   2091  100%  0.0   93.77  1458       AY631946.1
Helicobacter sp. MIT 95-234-6 16S ribosomal RNA gene, partial...  Helicobacter... NA              152487     2091   2091  100%  0.0   93.78  1444       AF336948.1
Helicobacter canadensis strain NCTC 13242 16S ribosomal RNA...    Helicobacter... NA              123841     2089   2089  100%  0.0   93.79  1453       KJ534295.1
Helicobacter rappini strain H 157 S 16S ribosomal RNA gene,...    Helicobacter... NA              95150      2089   2089  100%  0.0   93.77  1456       AY034821.1
Helicobacter sp. BTP3S 16S ribosomal RNA gene, partial sequence   Helicobacter... NA              281983     2089   2089  100%  0.0   93.79  1453       AY554137.1
Helicobacter canis strain Lausanne 16S ribosomal RNA gene,...     Helicobacter... NA              29419      2087   2087  100%  0.0   93.71  1465       DQ412573.1
Helicobacter rappini strain H 95 S 16S ribosomal RNA gene,...     Helicobacter... NA              95150      2087   2087  100%  0.0   93.77  1455       AY034820.1
Helicobacter sp. MIT 13-10041 16S ribosomal RNA gene, partial...  Helicobacter... NA              1680554    2085   2085  100%  0.0   93.71  1473       KT159710.1
Helicobacter sp. strain XJK30-2 16S ribosomal RNA gene, partia... Helicobacter... NA              218        2085   2085  100%  0.0   93.71  1474       OP278857.1
Helicobacter canis strain ATCC 51401 16S ribosomal RNA, partia... Helicobacter... NA              29419      2085   2085  100%  0.0   93.70  1473       NR_043052.1
Helicobacter sp. LNB2F 16S ribosomal RNA gene, partial sequence   Helicobacter... NA              281985     2085   2085  100%  0.0   93.78  1452       AY554139.1
Helicobacter sp. 'B10B Seymour' 16S ribosomal RNA gene, partia... Helicobacter... NA              279277     2084   2084  99%   0.0   93.71  1480       M88139.3
Helicobacter sp. BTP1S 16S ribosomal RNA gene, partial sequence   Helicobacter... NA              281981     2084   2084  100%  0.0   93.72  1480       AY554135.1
Helicobacter hepaticus strain Hh-2 16S ribosomal RNA, partial...  Helicobacter... NA              32025      2082   2082  100%  0.0   93.69  1470       NR_025952.1
Helicobacter hepaticus strain MIT 96-284 16S ribosomal RNA gen... Helicobacter... NA              32025      2082   2082  100%  0.0   93.69  1442       AY631953.1
Flexispira rappini 16S ribosomal RNA gene, partial sequence       Flexispira r... NA              2354       2082   2082  99%   0.0   93.70  1449       AF497807.1
Helicobacter rappini strain 2 SU 16S ribosomal RNA gene, parti... Helicobacter... NA              95150      2082   2082  100%  0.0   93.63  1456       AY034818.1
Helicobacter hepaticus ATCC 51449, complete genome                Helicobacter... NA              235279     2082   2082  100%  0.0   93.69  1799146    AE017125.1
Helicobacter hepaticus 16S ribosomal RNA (16S rRNA) gene          Helicobacter... NA              32025      2082   2082  100%  0.0   93.69  1457       L39122.1
Flexispira rappini strain 204-2 16S ribosomal RNA gene, partia... Flexispira r... NA              2354       2080   2080  100%  0.0   93.64  1457       AY192527.1
Flexispira rappini 16S ribosomal RNA gene, complete sequence      Flexispira r... NA              2354       2080   2080  100%  0.0   93.64  1487       AF034135.1
Helicobacter sp. strain HEX9 16S ribosomal RNA gene, partial...   Helicobacter... NA              218        2078   2078  99%   0.0   93.63  1496       OQ181203.1
Helicobacter sp. strain HAX4 16S ribosomal RNA gene, partial...   Helicobacter... NA              218        2078   2078  99%   0.0   93.64  1496       OQ181199.1
Helicobacter canadensis strain L222 16S ribosomal RNA gene,...    Helicobacter... NA              123841     2078   2078  99%   0.0   93.75  1403       DQ438119.1
Helicobacter sp. MIT 01-5529A 16S ribosomal RNA gene, partial...  Helicobacter... NA              222133     2078   2078  100%  0.0   93.63  1453       AY203898.1
Helicobacter hepaticus strain ATCC 51448 16S ribosomal RNA,...    Helicobacter... NA              32025      2078   2078  99%   0.0   93.68  1445       NR_114584.1
Helicobacter sp. 'Flexispira taxon 4' 16S ribosomal RNA gene,...  Helicobacter... NA              50960      2078   2078  100%  0.0   93.63  1438       AF225548.1
Helicobacter sp. PAGU 1991 PAGU 1750 gene for 16S rRNA, partia... Helicobacter... NA              2875691    2076   2076  100%  0.0   93.64  1417       LC650921.1
Helicobacter sp. MIT 11-6608 16S ribosomal RNA gene, partial...   Helicobacter... NA              1332284    2076   2076  99%   0.0   93.63  1472       KC894697.1
Uncultured organism clone ELU0021-T97-S-NI_000164 small subuni... uncultured o... NA              155900     2076   2076  100%  0.0   93.62  1413       HQ746686.1
Helicobacter himalayensis strain YS1, complete genome             Helicobacter... NA              1591088    2074   4149  100%  0.0   93.57  1829936    CP014991.1
Helicobacter himalayensis strain 80(YS1) 16S ribosomal RNA,...    Helicobacter... NA              1591088    2074   2074  100%  0.0   93.57  1487       NR_135861.1
Helicobacter cinaedi 94105 DNA, complete genome                   Helicobacter... NA              213        2074   4149  100%  0.0   93.57  1771717    AP025204.1
Helicobacter callitrichis strain R-204 16S ribosomal RNA gene,... Flexispira r... NA              2354       2074   2074  100%  0.0   93.57  1457       AY192526.1
Helicobacter cinaedi strain MIT 01-5002 16S ribosomal RNA gene... Helicobacter... NA              213        2074   2074  100%  0.0   93.57  1473       AY631947.1
Flexispira sp. CDC-H69 16S ribosomal RNA gene, partial sequence   Flexispira s... NA              98428      2074   2074  100%  0.0   93.57  1488       AF118807.1
Helicobacter valdiviensis strain SuratThani2016 16S ribosomal...  Helicobacter... NA              1458358    2073   2073  99%   0.0   93.75  1400       KX503247.1
Helicobacter pullorum strain 3758-94 16S ribosomal RNA gene,...   Helicobacter... NA              35818      2073   2073  100%  0.0   93.57  1462       KJ534305.1
Uncultured organism clone ELU0008-T58-S-NI_000407 small subuni... uncultured o... NA              155900     2073   2073  99%   0.0   93.61  1412       HQ740314.1
Helicobacter sp. MIT 02-5519C 16S ribosomal RNA gene, partial...  Helicobacter... NA              222135     2073   2073  100%  0.0   93.57  1453       AY203900.1
Helicobacter jaachi strain MIT 09-6948 16S ribosomal RNA gene,... Helicobacter... NA              1677920    2071   2071  95%   0.0   94.88  1427       KP701327.1
Helicobacter sp. PAGU 1991 PAGU 1991 gene for 16S rRNA, partia... Helicobacter... NA              2875691    2071   2071  100%  0.0   93.57  1417       LC650920.1
Uncultured organism clone ELU0021-T97-S-NI_000154 small subuni... uncultured o... NA              155900     2071   2071  99%   0.0   93.60  1413       HQ746676.1
Uncultured organism clone ELU0008-T58-S-NI_000374 small subuni... uncultured o... NA              155900     2071   2071  99%   0.0   93.71  1417       HQ740281.1
Helicobacter cinaedi 16S ribosomal RNA gene, partial sequence     Helicobacter... NA              213        2071   2071  99%   0.0   93.56  1448       AF497810.1
Helicobacter cf. pullorum strain W.Tee-Cro 16S ribosomal RNA...   Helicobacter... NA              172453     2071   2071  100%  0.0   93.57  1479       AF334681.1
Helicobacter sp. MIT 99-5507 16S ribosomal RNA gene, partial...   Helicobacter... NA              152489     2071   2071  99%   0.0   93.56  1490       AF333340.1
Helicobacter sp. CNRCH-2013/518 16S ribosomal RNA gene, partia... Helicobacter... NA              1501965    2069   2069  100%  0.0   93.51  1458       KJ534294.1
Helicobacter cinaedi PAGU611 DNA, complete genome                 Helicobacter... NA              1172562    2069   4138  100%  0.0   93.50  2078348    AP012344.1
Helicobacter cinaedi gene for 16S rRNA, partial sequence,...      Helicobacter... NA              1172562    2069   2069  100%  0.0   93.50  1429       AB275318.1
Helicobacter rappini W.Tee-Yu 16S ribosomal RNA gene, partial...  Helicobacter... NA              95150      2069   2069  100%  0.0   93.51  1486       AF286053.1
Flexispira rappini 16S ribosomal RNA gene, partial sequence       Flexispira r... NA              2354       2069   2069  100%  0.0   93.50  1451       AF118017.1
Helicobacter sp. MIT 17-337 16S ribosomal RNA gene, partial...    Helicobacter... NA              2040648    2067   2067  100%  0.0   93.50  1504       MH726196.1
Uncultured Helicobacter sp. clone F37_1 16S ribosomal RNA gene... uncultured H... NA              175537     2067   2067  99%   0.0   93.49  1458       PP463019.1
Helicobacter didelphidarum strain MIT 17-337 16S ribosomal RNA... Helicobacter... NA              2040648    2067   2067  100%  0.0   93.50  1497       NR_180169.1
Helicobacter colisuis 16S ribosomal RNA gene, partial sequence    Helicobacter... NA              2949739    2067   2067  100%  0.0   93.50  1494       OM338647.1
Helicobacter sp. MIT 03-7674c 16S ribosomal RNA gene, partial...  Helicobacter... NA              398628     2067   2067  100%  0.0   93.50  1453       DQ846679.1
Helicobacter canadensis strain L178 16S ribosomal RNA gene,...    Helicobacter... NA              123841     2067   2067  100%  0.0   93.50  1447       DQ438114.1
Helicobacter canadensis strain L172 16S ribosomal RNA gene,...    Helicobacter... NA              123841     2067   2067  100%  0.0   93.50  1446       DQ438113.1
Helicobacter canadensis isolate Gotland B75 16S ribosomal RNA...  Helicobacter... NA              123841     2067   2067  99%   0.0   93.66  1394       AY323505.1
Helicobacter pullorum NCTC 12824 strain ATCC 51801 16S ribosom... Helicobacter... NA              552535     2067   2067  100%  0.0   93.50  1473       NR_043053.1
Helicobacter marmotae strain MIT 98-6070 16S ribosomal RNA,...    Helicobacter... NA              152490     2067   2067  100%  0.0   93.50  1473       NR_041825.1
Helicobacter cinaedi strain ADN 0413 16S ribosomal RNA gene,...   Helicobacter... NA              1869279    2067   2067  99%   0.0   93.72  1402       AF207737.1
Helicobacter sp. WB2F 16S ribosomal RNA gene, partial sequence    Helicobacter... NA              281987     2067   2067  100%  0.0   93.49  1484       AY554141.1
Helicobacter sp. WB1F 16S ribosomal RNA gene, partial sequence    Helicobacter... NA              281986     2067   2067  100%  0.0   93.50  1474       AY554140.1
Helicobacter didelphidarum strain MIT 16-1508 16S ribosomal RN... Helicobacter... NA              2040648    2065   2065  100%  0.0   93.50  1457       MN630626.1
Helicobacter sp. MIT 15-1451 16S ribosomal RNA gene, partial...   Helicobacter... NA              2293435    2065   2065  99%   0.0   93.49  1504       MH726195.1
Helicobacter monodelphidis strain MIT 15-1451 16S ribosomal RN... Helicobacter... NA              2293435    2065   2065  99%   0.0   93.49  1497       NR_180168.1
Uncultured organism clone ELU0021-T97-S-NI_000090 small subuni... uncultured o... NA              155900     2065   2065  99%   0.0   93.64  1408       HQ746612.1
Helicobacter equorum gene for 16S ribosomal RNA, partial...       Helicobacter... NA              361872     2065   2065  100%  0.0   93.50  1445       AB571486.1
Helicobacter hepaticus strain MIT 96-1809 16S ribosomal RNA...    Helicobacter... NA              32025      2065   2065  100%  0.0   93.47  1441       AY631952.1
Helicobacter canicola gene for 16S ribosomal RNA, partial...      Helicobacter... NA              1869279    2063   2063  99%   0.0   93.60  1396       LC102853.1
Helicobacter canis strain MIT 12-7709 16S ribosomal RNA gene,...  Helicobacter... NA              29419      2063   2063  100%  0.0   93.43  1459       KC878293.1
Helicobacter cinaedi strain 11-30771 16S ribosomal RNA gene,...   Helicobacter... NA              213        2063   2063  100%  0.0   93.42  1451       KC616311.1
Helicobacter cinaedi strain 09-31248 16S ribosomal RNA gene,...   Helicobacter... NA              213        2063   2063  100%  0.0   93.42  1451       KC616310.1
Helicobacter sp. 'Flexispira taxon 8' 16S ribosomal RNA gene,...  Helicobacter... NA              613026     2063   2063  100%  0.0   93.42  1440       M88138.3


Alignments:

>Uncultured Helicobacter sp. clone R16-645 16S ribosomal RNA gene, partial sequence
Sequence ID: MT758473.1 Length: 1391
Range 1: 1 to 1391

Score:2569 bits(1391), Expect:0.0,
Identities:1391/1391(100%),  Gaps:0/1391(0%), Strand: Plus/Plus

Query  1     AGTGAACGCGGCGGCGTGCCTAATACATGCAAGTCGAACGAAGTTATAAGGGCTTGCCTT  60
             ||||||||||||||||||||||||||||||||||||||||||||||||||||||||||||
Sbjct  1     AGTGAACGCGGCGGCGTGCCTAATACATGCAAGTCGAACGAAGTTATAAGGGCTTGCCTT  60

Query  61    TATAACTTAGTGGCGCACGGGTGAGTAATGCATAGGTAACATGCCTCATAGTCTGGGATA  120
             ||||||||||||||||||||||||||||||||||||||||||||||||||||||||||||
Sbjct  61    TATAACTTAGTGGCGCACGGGTGAGTAATGCATAGGTAACATGCCTCATAGTCTGGGATA  120

Query  121   GCCACTGGAAACGGTGATTAATACTAGATACTCCTTACGAGGGAAAGAATTTCGCTATGA  180
             ||||||||||||||||||||||||||||||||||||||||||||||||||||||||||||
Sbjct  121   GCCACTGGAAACGGTGATTAATACTAGATACTCCTTACGAGGGAAAGAATTTCGCTATGA  180

Query  181   GATTGGCCTATGTCCCATCAGCTTGTTGGTAAGGTAATGGCTTACCAAGGCTATGACGGG  240
             ||||||||||||||||||||||||||||||||||||||||||||||||||||||||||||
Sbjct  181   GATTGGCCTATGTCCCATCAGCTTGTTGGTAAGGTAATGGCTTACCAAGGCTATGACGGG  240

Query  241   TATCCGGCCTGAGAGGGTGAACGGACACACTGGAACTGAGACACGGTCCAGACTCCTACG  300
             ||||||||||||||||||||||||||||||||||||||||||||||||||||||||||||
Sbjct  241   TATCCGGCCTGAGAGGGTGAACGGACACACTGGAACTGAGACACGGTCCAGACTCCTACG  300

Query  301   GGAGGCAGCAGTAGGGAATATTGCTCAATGGGGGAAACCCTGAAGCAGCAACGCCGCGTG  360
             ||||||||||||||||||||||||||||||||||||||||||||||||||||||||||||
Sbjct  301   GGAGGCAGCAGTAGGGAATATTGCTCAATGGGGGAAACCCTGAAGCAGCAACGCCGCGTG  360

Query  361   GAGGATGAAGGTTTTAGGATTGTAAACTCCTTTTCTAAGAGAAGATTATGACGGTATCTT  420
             ||||||||||||||||||||||||||||||||||||||||||||||||||||||||||||
Sbjct  361   GAGGATGAAGGTTTTAGGATTGTAAACTCCTTTTCTAAGAGAAGATTATGACGGTATCTT  420

Query  421   AGGAATAAGCACCGGCTAACTCCGTGCCAGCAGCCGCGGTAATACGGAGGGTGCAAGCGT  480
             ||||||||||||||||||||||||||||||||||||||||||||||||||||||||||||
Sbjct  421   AGGAATAAGCACCGGCTAACTCCGTGCCAGCAGCCGCGGTAATACGGAGGGTGCAAGCGT  480

Query  481   TACTCGGAATCACTGGGCGTAAAGAGCGCGTAGGCGGAATAACAAGTCAGATGTGAAATC  540
             ||||||||||||||||||||||||||||||||||||||||||||||||||||||||||||
Sbjct  481   TACTCGGAATCACTGGGCGTAAAGAGCGCGTAGGCGGAATAACAAGTCAGATGTGAAATC  540

Query  541   CTGTAGCTTAACTACAGAACTGCATTTGAAACTGTTGTTCTAGAGTGTGGGAGAGGTAGG  600
             ||||||||||||||||||||||||||||||||||||||||||||||||||||||||||||
Sbjct  541   CTGTAGCTTAACTACAGAACTGCATTTGAAACTGTTGTTCTAGAGTGTGGGAGAGGTAGG  600

Query  601   TGGAATTCTTGGTGTAGGGGTAAAATCCGTAGAGATCAAGAGGAATACTCATTGCGAAGG  660
             ||||||||||||||||||||||||||||||||||||||||||||||||||||||||||||
Sbjct  601   TGGAATTCTTGGTGTAGGGGTAAAATCCGTAGAGATCAAGAGGAATACTCATTGCGAAGG  660

Query  661   CGACCTACTGGAACATTACTGACGCTGATGCGCGAAAGCGTGGGGAGCAAACAGGATTAG  720
             ||||||||||||||||||||||||||||||||||||||||||||||||||||||||||||
Sbjct  661   CGACCTACTGGAACATTACTGACGCTGATGCGCGAAAGCGTGGGGAGCAAACAGGATTAG  720

Query  721   ATACCCTGGTAGTCCACGCCCTAAACTATGGATGCTAGTTGTTGCCTTGCTAGTCAAGGC  780
             ||||||||||||||||||||||||||||||||||||||||||||||||||||||||||||
Sbjct  721   ATACCCTGGTAGTCCACGCCCTAAACTATGGATGCTAGTTGTTGCCTTGCTAGTCAAGGC  780

Query  781   AGTAATGCAGCTAACGCATTAAGCATCCCGCCTGGGGAGTACGGTCGCAAGATTAAAACT  840
             ||||||||||||||||||||||||||||||||||||||||||||||||||||||||||||
Sbjct  781   AGTAATGCAGCTAACGCATTAAGCATCCCGCCTGGGGAGTACGGTCGCAAGATTAAAACT  840

Query  841   CAAAGGAATAGACGGGGACCCGCACAAGCGGTGGAGCATGTGGTTTAATTCGAAGATACG  900
             ||||||||||||||||||||||||||||||||||||||||||||||||||||||||||||
Sbjct  841   CAAAGGAATAGACGGGGACCCGCACAAGCGGTGGAGCATGTGGTTTAATTCGAAGATACG  900

Query  901   CGAAGAACCTTACCTAGGCTTGACATTGATAGAATCCGCTAGAGATAGTGGAGTGCCACG  960
             ||||||||||||||||||||||||||||||||||||||||||||||||||||||||||||
Sbjct  901   CGAAGAACCTTACCTAGGCTTGACATTGATAGAATCCGCTAGAGATAGTGGAGTGCCACG  960

Query  961   CAAGTGGAGCTTGAAAACAGGTGCTGCACGGCTGTCGTCAGCTCGTGTCGTGAGATGTTG  1020
             ||||||||||||||||||||||||||||||||||||||||||||||||||||||||||||
Sbjct  961   CAAGTGGAGCTTGAAAACAGGTGCTGCACGGCTGTCGTCAGCTCGTGTCGTGAGATGTTG  1020

Query  1021  GGTTAAGTCCCGCAACGAGCGCAACCCTTATCCTTAGTTGCTAGCAGTTAGGCTGAGCAC  1080
             ||||||||||||||||||||||||||||||||||||||||||||||||||||||||||||
Sbjct  1021  GGTTAAGTCCCGCAACGAGCGCAACCCTTATCCTTAGTTGCTAGCAGTTAGGCTGAGCAC  1080

Query  1081  TCTAAGGAGACTGCCTTCGTAAGAAGGAGGAAGGCGAGGACGACGTCAAGTCATCATGGC  1140
             ||||||||||||||||||||||||||||||||||||||||||||||||||||||||||||
Sbjct  1081  TCTAAGGAGACTGCCTTCGTAAGAAGGAGGAAGGCGAGGACGACGTCAAGTCATCATGGC  1140

Query  1141  CCTTACGCCTAGGGCTACACACGTGCTACAATGGTTGATACAAAGAGATGCAATACTGCG  1200
             ||||||||||||||||||||||||||||||||||||||||||||||||||||||||||||
Sbjct  1141  CCTTACGCCTAGGGCTACACACGTGCTACAATGGTTGATACAAAGAGATGCAATACTGCG  1200

Query  1201  AAGTGGAGCCAATCTCAAAAATCAATCTCAGTTCGGATTGTAGGCTGCAACTCGCCTACA  1260
             ||||||||||||||||||||||||||||||||||||||||||||||||||||||||||||
Sbjct  1201  AAGTGGAGCCAATCTCAAAAATCAATCTCAGTTCGGATTGTAGGCTGCAACTCGCCTACA  1260

Query  1261  TGAAGCTGGAATCGCTAGTAATCGTAAATCAGCAATGTTACGGTGAATACGTTCCCGGGT  1320
             ||||||||||||||||||||||||||||||||||||||||||||||||||||||||||||
Sbjct  1261  TGAAGCTGGAATCGCTAGTAATCGTAAATCAGCAATGTTACGGTGAATACGTTCCCGGGT  1320

Query  1321  CTTGTACTCACCGCCCGTCACACCATGGGAGTTGTATTCGCCTTAAGTCGGAATACTAAA  1380
             ||||||||||||||||||||||||||||||||||||||||||||||||||||||||||||
Sbjct  1321  CTTGTACTCACCGCCCGTCACACCATGGGAGTTGTATTCGCCTTAAGTCGGAATACTAAA  1380

Query  1381  TTAGTTACCGC  1391
             |||||||||||
Sbjct  1381  TTAGTTACCGC  1391


>Helicobacter sp. 11S02629-2 16S ribosomal RNA gene, partial sequence
Sequence ID: KJ081205.1 Length: 1391
Range 1: 1 to 1387

Score:2200 bits(1191), Expect:0.0,
Identities:1327/1392(95%),  Gaps:11/1392(0%), Strand: Plus/Plus

Query  6     ACGC-GGCGGCGTGCCTAATACATGCAAGTCGAACGA---AG-TTATAAGGGCTTGCCTT  60
             |||| ||||||||||||||||||||||||||||||||   || || | |  ||||||
Sbjct  1     ACGCTGGCGGCGTGCCTAATACATGCAAGTCGAACGATGAAGCTTCT-A--GCTTGCTAG  57

Query  61    TA-TAACTTAGTGGCGCACGGGTGAGTAATGCATAGGTAACATGCCTCATAGTCTGGGAT  119
              | |   |||||||||||||||||||||||||||||||||||||||   |||||||||||
Sbjct  58    AAGTGGATTAGTGGCGCACGGGTGAGTAATGCATAGGTAACATGCCCTTTAGTCTGGGAT  117

Query  120   AGCCACTGGAAACGGTGATTAATACTAGATACTCCTTACGAGGGAAAGAATTTCGCTATG  179
             || |||| |||| ||||| |||||||||||||||| |||| |||||||||||||||||
Sbjct  118   AGTCACTAGAAATGGTGAATAATACTAGATACTCCCTACGGGGGAAAGAATTTCGCTAAA  177

Query  180   AGATTGGCCTATGTCCCATCAGCTTGTTGGTAAGGTAATGGCTTACCAAGGCTATGACGG  239
              |||||||||||||||||||||||||||||||||||||||||||||||||||||||||||
Sbjct  178   GGATTGGCCTATGTCCCATCAGCTTGTTGGTAAGGTAATGGCTTACCAAGGCTATGACGG  237

Query  240   GTATCCGGCCTGAGAGGGTGAACGGACACACTGGAACTGAGACACGGTCCAGACTCCTAC  299
             ||||||||||||||||||||||||||||||||||||||||||||||||||||||||||||
Sbjct  238   GTATCCGGCCTGAGAGGGTGAACGGACACACTGGAACTGAGACACGGTCCAGACTCCTAC  297

Query  300   GGGAGGCAGCAGTAGGGAATATTGCTCAATGGGGGAAACCCTGAAGCAGCAACGCCGCGT  359
             ||||||||||||||||||||||||||||||||||||||||||||||||||||||||||||
Sbjct  298   GGGAGGCAGCAGTAGGGAATATTGCTCAATGGGGGAAACCCTGAAGCAGCAACGCCGCGT  357

Query  360   GGAGGATGAAGGTTTTAGGATTGTAAACTCCTTTTCTAAGAGAAGATTATGACGGTATCT  419
             ||||||||||||||||||||||||||||||||||||||||||||||||||||||||||||
Sbjct  358   GGAGGATGAAGGTTTTAGGATTGTAAACTCCTTTTCTAAGAGAAGATTATGACGGTATCT  417

Query  420   TAGGAATAAGCACCGGCTAACTCCGTGCCAGCAGCCGCGGTAATACGGAGGGTGCAAGCG  479
             ||||||||||||||||||||||||||||||||||||||||||||||||||||||||||||
Sbjct  418   TAGGAATAAGCACCGGCTAACTCCGTGCCAGCAGCCGCGGTAATACGGAGGGTGCAAGCG  477

Query  480   TTACTCGGAATCACTGGGCGTAAAGAGCGCGTAGGCGGAATAACAAGTCAGATGTGAAAT  539
             |||||||||||||||||||||||||||||||||||||| |||| ||||||||||||||||
Sbjct  478   TTACTCGGAATCACTGGGCGTAAAGAGCGCGTAGGCGGGATAATAAGTCAGATGTGAAAT  537

Query  540   CCTGTAGCTTAACTACAGAACTGCATTTGAAACTGTTGTTCTAGAGTGTGGGAGAGGTAG  599
             ||||||||||||||||||||||||||||||||||||| ||||||||| ||||||||||||
Sbjct  538   CCTGTAGCTTAACTACAGAACTGCATTTGAAACTGTTATTCTAGAGTATGGGAGAGGTAG  597

Query  600   GTGGAATTCTTGGTGTAGGGGTAAAATCCGTAGAGATCAAGAGGAATACTCATTGCGAAG  659
             ||||||||||||||||||||||||||||||||||||||||||||||||||||||||||||
Sbjct  598   GTGGAATTCTTGGTGTAGGGGTAAAATCCGTAGAGATCAAGAGGAATACTCATTGCGAAG  657

Query  660   GCGACCTACTGGAACATTACTGACGCTGATGCGCGAAAGCGTGGGGAGCAAACAGGATTA  719
             ||||||||||||||||||||||||||||||||||||||||||||||||||||||||||||
Sbjct  658   GCGACCTACTGGAACATTACTGACGCTGATGCGCGAAAGCGTGGGGAGCAAACAGGATTA  717

Query  720   GATACCCTGGTAGTCCACGCCCTAAACTATGGATGCTAGTTGTTGCCTTGCTAGTCAAGG  779
             |||||||||||||||||||||||||||||||||||||||||||||    |||||||
Sbjct  718   GATACCCTGGTAGTCCACGCCCTAAACTATGGATGCTAGTTGTTGGGGAGCTAGTCTCTC  777

Query  780   CAGTAATGCAGCTAACGCATTAAGCATCCCGCCTGGGGAGTACGGTCGCAAGATTAAAAC  839
             ||||||||||||||||||||||||||||||||||||||||||||||||||||||||||||
Sbjct  778   CAGTAATGCAGCTAACGCATTAAGCATCCCGCCTGGGGAGTACGGTCGCAAGATTAAAAC  837

Query  840   TCAAAGGAATAGACGGGGACCCGCACAAGCGGTGGAGCATGTGGTTTAATTCGAAGATAC  899
             ||||||||||||||||||||||||||||||||||||||||||||||||||||||||||||
Sbjct  838   TCAAAGGAATAGACGGGGACCCGCACAAGCGGTGGAGCATGTGGTTTAATTCGAAGATAC  897

Query  900   GCGAAGAACCTTACCTAGGCTTGACATTGATAGAATCCGCTAGAGATAGTGGAGTGCCAC  959
              |||||||||||||||||||||||||||||||||||| ||||||||||| ||||||||
Sbjct  898   ACGAAGAACCTTACCTAGGCTTGACATTGATAGAATCTGCTAGAGATAGCGGAGTGCCCT  957

Query  960   GCAAGTGGAGCTTGAAAACAGGTGCTGCACGGCTGTCGTCAGCTCGTGTCGTGAGATGTT  1019
              |  | ||||||||||||||||||||||||||||||||||||||||||||||||||||||
Sbjct  958   TCG-G-GGAGCTTGAAAACAGGTGCTGCACGGCTGTCGTCAGCTCGTGTCGTGAGATGTT  1015

Query  1020  GGGTTAAGTCCCGCAACGAGCGCAACCCTTATCCTTAGTTGCTAGCAGTTAGGCTGAGCA  1079
             |||||||||||||||||||||||||||||  ||||||||||||||||||| |||||||||
Sbjct  1016  GGGTTAAGTCCCGCAACGAGCGCAACCCTCGTCCTTAGTTGCTAGCAGTTCGGCTGAGCA  1075

Query  1080  CTCTAAGGAGACTGCCTTCGTAAGAAGGAGGAAGGCGAGGACGACGTCAAGTCATCATGG  1139
             || ||||||||||||||| |||||||||||||||| ||||||||||||||||||||||||
Sbjct  1076  CTATAAGGAGACTGCCTTTGTAAGAAGGAGGAAGGTGAGGACGACGTCAAGTCATCATGG  1135

Query  1140  CCCTTACGCCTAGGGCTACACACGTGCTACAATGGTTGATACAAAGAGATGCAATACTGC  1199
             ||||||||||||||||||||||||||||||||||||||||||||||||||||||||| ||
Sbjct  1136  CCCTTACGCCTAGGGCTACACACGTGCTACAATGGTTGATACAAAGAGATGCAATACCGC  1195

Query  1200  GAAGTGGAGCCAATCTCAAAAATCAATCTCAGTTCGGATTGTAGGCTGCAACTCGCCTAC  1259
             || ||||||| |||||||||||| ||||||||||||||||| |||||||||||||||| |
Sbjct  1196  GAGGTGGAGCAAATCTCAAAAATTAATCTCAGTTCGGATTGAAGGCTGCAACTCGCCTTC  1255

Query  1260  ATGAAGCTGGAATCGCTAGTAATCGTAAATCAGCAATGTTACGGTGAATACGTTCCCGGG  1319
             ||||||||||||||||||||||||||| |||||| ||| |||||||||||||||||||||
Sbjct  1256  ATGAAGCTGGAATCGCTAGTAATCGTAGATCAGCGATGCTACGGTGAATACGTTCCCGGG  1315

Query  1320  TCTTGTACTCACCGCCCGTCACACCATGGGAGTTGTATTCGCCTTAAGTCGGAATACTAA  1379
             ||||||||||||||||||||||||||||||||||||||||||||||||||||||||||||
Sbjct  1316  TCTTGTACTCACCGCCCGTCACACCATGGGAGTTGTATTCGCCTTAAGTCGGAATACTAA  1375

Query  1380  ATTAGTTACCGC  1391
             ||||||||||||
Sbjct  1376  ATTAGTTACCGC  1387


>Helicobacter sp. strain MIT 16-1353 16S ribosomal RNA gene, partial sequence
Sequence ID: MW147609.1 Length: 1504
Range 1: 31 to 1423

Score:2170 bits(1175), Expect:0.0,
Identities:1321/1393(95%),  Gaps:3/1393(0%), Strand: Plus/Plus

Query  1     AGTGAACGC-GGCGGCGTGCCTAATACATGCAAGTCGAACGAAGTTAT-AAGGGCTTGCC  58
             ||||||||| |||||||||||||||||||||||||||||||| |  ||     ||||||
Sbjct  31    AGTGAACGCTGGCGGCGTGCCTAATACATGCAAGTCGAACGATGAAATTTCTAGCTTGCT  90

Query  59    TTTA-TAACTTAGTGGCGCACGGGTGAGTAATGCATAGGTAACATGCCTCATAGTCTGGG  117
                | |   ||||||||||||||||||||||||||||||| |  ||||   |||||| ||
Sbjct  91    AGAAGTGGATTAGTGGCGCACGGGTGAGTAATGCATAGGTTATGTGCCCTTTAGTCTAGG  150

Query  118   ATAGCCACTGGAAACGGTGATTAATACTAGATACTCCTTACGAGGGAAAGAATTTCGCTA  177
             |||||||||||||||||||||||||||| |||||||||||||||||||||||||||||||
Sbjct  151   ATAGCCACTGGAAACGGTGATTAATACTGGATACTCCTTACGAGGGAAAGAATTTCGCTA  210

Query  178   TGAGATTGGCCTATGTCCCATCAGCTTGTTGGTAAGGTAATGGCTTACCAAGGCTATGAC  237
                |||  |||||||||| |||||||||||||| ||||||||||| ||||||||||||||
Sbjct  211   AAGGATCAGCCTATGTCCTATCAGCTTGTTGGTGAGGTAATGGCTCACCAAGGCTATGAC  270

Query  238   GGGTATCCGGCCTGAGAGGGTGAACGGACACACTGGAACTGAGACACGGTCCAGACTCCT  297
             ||||||||||||||||||||||||||||||||||||||||||||||||||||||||||||
Sbjct  271   GGGTATCCGGCCTGAGAGGGTGAACGGACACACTGGAACTGAGACACGGTCCAGACTCCT  330

Query  298   ACGGGAGGCAGCAGTAGGGAATATTGCTCAATGGGGGAAACCCTGAAGCAGCAACGCCGC  357
             ||||||||||||||||||||||||||||||||||||||||||||||||||||||||||||
Sbjct  331   ACGGGAGGCAGCAGTAGGGAATATTGCTCAATGGGGGAAACCCTGAAGCAGCAACGCCGC  390

Query  358   GTGGAGGATGAAGGTTTTAGGATTGTAAACTCCTTTTCTAAGAGAAGATTATGACGGTAT  417
             ||||||||||||||||||||||||||||||||||||||||||||||||||||||||||||
Sbjct  391   GTGGAGGATGAAGGTTTTAGGATTGTAAACTCCTTTTCTAAGAGAAGATTATGACGGTAT  450

Query  418   CTTAGGAATAAGCACCGGCTAACTCCGTGCCAGCAGCCGCGGTAATACGGAGGGTGCAAG  477
             ||||||||||||||||||||||||||||||||||||||||||||||||||||||||||||
Sbjct  451   CTTAGGAATAAGCACCGGCTAACTCCGTGCCAGCAGCCGCGGTAATACGGAGGGTGCAAG  510

Query  478   CGTTACTCGGAATCACTGGGCGTAAAGAGCGCGTAGGCGGAATAACAAGTCAGATGTGAA  537
             ||||||||||||||||||||||||||||||||||||||||||||| ||||||||||||||
Sbjct  511   CGTTACTCGGAATCACTGGGCGTAAAGAGCGCGTAGGCGGAATAATAAGTCAGATGTGAA  570

Query  538   ATCCTGTAGCTTAACTACAGAACTGCATTTGAAACTGTTGTTCTAGAGTGTGGGAGAGGT  597
             ||||| ||||||||||| |||| ||||||||||||| |||||||||||| ||||||||||
Sbjct  571   ATCCTATAGCTTAACTATAGAATTGCATTTGAAACTATTGTTCTAGAGTATGGGAGAGGT  630

Query  598   AGGTGGAATTCTTGGTGTAGGGGTAAAATCCGTAGAGATCAAGAGGAATACTCATTGCGA  657
             | ||||||||||||||||||||||||||||||||||||||||||||||||||||||||||
Sbjct  631   AAGTGGAATTCTTGGTGTAGGGGTAAAATCCGTAGAGATCAAGAGGAATACTCATTGCGA  690

Query  658   AGGCGACCTACTGGAACATTACTGACGCTGATGCGCGAAAGCGTGGGGAGCAAACAGGAT  717
             ||||||| ||||||||||||||||||||||||||||||||||||||||||||||||||||
Sbjct  691   AGGCGACTTACTGGAACATTACTGACGCTGATGCGCGAAAGCGTGGGGAGCAAACAGGAT  750

Query  718   TAGATACCCTGGTAGTCCACGCCCTAAACTATGGATGCTAGTTGTTGCCTTGCTAGTCAA  777
             ||||||||||||||||||||||||||||||||| |||||||||||||||||||| ||||
Sbjct  751   TAGATACCCTGGTAGTCCACGCCCTAAACTATGAATGCTAGTTGTTGCCTTGCTTGTCAG  810

Query  778   GGCAGTAATGCAGCTAACGCATTAAGCATCCCGCCTGGGGAGTACGGTCGCAAGATTAAA  837
             ||||||||||||||||||||||||||||| ||||||||||||||||||||||||||||||
Sbjct  811   GGCAGTAATGCAGCTAACGCATTAAGCATTCCGCCTGGGGAGTACGGTCGCAAGATTAAA  870

Query  838   ACTCAAAGGAATAGACGGGGACCCGCACAAGCGGTGGAGCATGTGGTTTAATTCGAAGAT  897
             ||||||||||||||||||||||||||||||||||||||||||||||||||||||||||||
Sbjct  871   ACTCAAAGGAATAGACGGGGACCCGCACAAGCGGTGGAGCATGTGGTTTAATTCGAAGAT  930

Query  898   ACGCGAAGAACCTTACCTAGGCTTGACATTGATAGAATCCGCTAGAGATAGTGGAGTGCC  957
             ||||||||||||||||||||||||||||||||||||||| ||||||||||| ||||||||
Sbjct  931   ACGCGAAGAACCTTACCTAGGCTTGACATTGATAGAATCTGCTAGAGATAGCGGAGTGCC  990

Query  958   ACGCAAGTGGAGCTTGAAAACAGGTGCTGCACGGCTGTCGTCAGCTCGTGTCGTGAGATG  1017
             ||||||||||||||||||||||||||||||||||||||||||||||||||||||||||||
Sbjct  991   ACGCAAGTGGAGCTTGAAAACAGGTGCTGCACGGCTGTCGTCAGCTCGTGTCGTGAGATG  1050

Query  1018  TTGGGTTAAGTCCCGCAACGAGCGCAACCCTTATCCTTAGTTGCTAGCAGTTAGGCTGAG  1077
             |||||||||||||||||||||||||||||||||||||||||||||||||||| |||||||
Sbjct  1051  TTGGGTTAAGTCCCGCAACGAGCGCAACCCTTATCCTTAGTTGCTAGCAGTTCGGCTGAG  1110

Query  1078  CACTCTAAGGAGACTGCCTTCGTAAGAAGGAGGAAGGCGAGGACGACGTCAAGTCATCAT  1137
             |||||||||||||||||||| |||| ||||||||||| ||||||||||||||||||||||
Sbjct  1111  CACTCTAAGGAGACTGCCTTTGTAAAAAGGAGGAAGGTGAGGACGACGTCAAGTCATCAT  1170

Query  1138  GGCCCTTACGCCTAGGGCTACACACGTGCTACAATGGTTGATACAAAGAGATGCAATACT  1197
             ||||||||||||||||||||||||||||||||||||||    ||||||||| |||||| |
Sbjct  1171  GGCCCTTACGCCTAGGGCTACACACGTGCTACAATGGTAAGCACAAAGAGAAGCAATATT  1230

Query  1198  GCGAAGTGGAGCCAATCTCAAAAATCAATCTCAGTTCGGATTGTAGGCTGCAACTCGCCT  1257
             ||||| |||||| |||||||||||   ||||||||||||||||||| |||||||||| ||
Sbjct  1231  GCGAAATGGAGCTAATCTCAAAAACTTATCTCAGTTCGGATTGTAGTCTGCAACTCGACT  1290

Query  1258  ACATGAAGCTGGAATCGCTAGTAATCGTAAATCAGCAATGTTACGGTGAATACGTTCCCG  1317
             |||||||||||||||||||||||||||| |||||||||||| ||||||||||||||||||
Sbjct  1291  ACATGAAGCTGGAATCGCTAGTAATCGTGAATCAGCAATGTCACGGTGAATACGTTCCCG  1350

Query  1318  GGTCTTGTACTCACCGCCCGTCACACCATGGGAGTTGTATTCGCCTTAAGTCGGAATACT  1377
             ||||||||||||||||||||||| |||||||||||||||||||||||||| ||| || ||
Sbjct  1351  GGTCTTGTACTCACCGCCCGTCAAACCATGGGAGTTGTATTCGCCTTAAGCCGGGATGCT  1410

Query  1378  AAATTAGTTACCG  1390
             ||| ||| |||||
Sbjct  1411  AAAATAGCTACCG  1423


>Helicobacter sp. 'pancake tortoise' 16S ribosomal RNA gene, partial sequence
Sequence ID: FJ667779.1 Length: 1370
Range 1: 46 to 1369

Score:2169 bits(1174), Expect:0.0,
Identities:1275/1325(96%),  Gaps:1/1325(0%), Strand: Plus/Plus

Query  67    TTAGTGGCGCACGGGTGAGTAATGCATAGGTAACATGCCTCATAGTCTGGGATAGCCACT  126
             |||||||||||||||||||||||||||||||||||||||   ||||||||||||||||||
Sbjct  46    TTAGTGGCGCACGGGTGAGTAATGCATAGGTAACATGCCCTTTAGTCTGGGATAGCCACT  105

Query  127   GGAAACGGTGATTAATACTAGATACTCCTTACGAGGGAAAGAATTTCGCTATGAGATTGG  186
              |||| |||||||||||||||||||| | |||| | |||||||||||||||   ||||||
Sbjct  106   AGAAATGGTGATTAATACTAGATACTTCCTACGGGAGAAAGAATTTCGCTAAAGGATTGG  165

Query  187   CCTATGTCCCATCAGCTTGTTGGTAAGGTAATGGCTTACCAAGGCTATGACGGGTATCCG  246
             ||||||||||||||||||||||||||||||||||||||||||||||||||||||||||||
Sbjct  166   CCTATGTCCCATCAGCTTGTTGGTAAGGTAATGGCTTACCAAGGCTATGACGGGTATCCG  225

Query  247   GCCTGAGAGGGTGAACGGACACACTGGAACTGAGACACGGTCCAGACTCCTACGGGAGGC  306
             ||||||||||||||||||||||||||||||||||||||||||||||||||||||||||||
Sbjct  226   GCCTGAGAGGGTGAACGGACACACTGGAACTGAGACACGGTCCAGACTCCTACGGGAGGC  285

Query  307   AGCAGTAGGGAATATTGCTCAATGGGGGAAACCCTGAAGCAGCAACGCCGCGTGGAGGAT  366
             ||||||||||||||||||||||||||||||||||||||||||||||||||||||||||||
Sbjct  286   AGCAGTAGGGAATATTGCTCAATGGGGGAAACCCTGAAGCAGCAACGCCGCGTGGAGGAT  345

Query  367   GAAGGTTTTAGGATTGTAAACTCCTTTTCTAAGAGAAGATTATGACGGTATCTTAGGAAT  426
             ||||||||||||||||||||||||||||||||||||||||||||||||||||||||||||
Sbjct  346   GAAGGTTTTAGGATTGTAAACTCCTTTTCTAAGAGAAGATTATGACGGTATCTTAGGAAT  405

Query  427   AAGCACCGGCTAACTCCGTGCCAGCAGCCGCGGTAATACGGAGGGTGCAAGCGTTACTCG  486
             ||||||||||||||||||||||||||||||||||||||||||||||||||||||||||||
Sbjct  406   AAGCACCGGCTAACTCCGTGCCAGCAGCCGCGGTAATACGGAGGGTGCAAGCGTTACTCG  465

Query  487   GAATCACTGGGCGTAAAGAGCGCGTAGGCGGAATAACAAGTCAGATGTGAAATCCTGTAG  546
             ||||||||||||||||||||||||||||||| |||| |||||||||||||||||||||||
Sbjct  466   GAATCACTGGGCGTAAAGAGCGCGTAGGCGGGATAATAAGTCAGATGTGAAATCCTGTAG  525

Query  547   CTTAACTACAGAACTGCATTTGAAACTGTTGTTCTAGAGTGTGGGAGAGGTAGGTGGAAT  606
             |||||||||||||||||||||||||||||| ||||||||| |||||||||||||||||||
Sbjct  526   CTTAACTACAGAACTGCATTTGAAACTGTTATTCTAGAGTATGGGAGAGGTAGGTGGAAT  585

Query  607   TCTTGGTGTAGGGGTAAAATCCGTAGAGATCAAGAGGAATACTCATTGCGAAGGCGACCT  666
             ||||||||||||||||||||||||||||||||||||||||||||||||||||||||||||
Sbjct  586   TCTTGGTGTAGGGGTAAAATCCGTAGAGATCAAGAGGAATACTCATTGCGAAGGCGACCT  645

Query  667   ACTGGAACATTACTGACGCTGATGCGCGAAAGCGTGGGGAGCAAACAGGATTAGATACCC  726
             ||||||||||||||||||||||||||||||||||||||||||||||||||||||||||||
Sbjct  646   ACTGGAACATTACTGACGCTGATGCGCGAAAGCGTGGGGAGCAAACAGGATTAGATACCC  705

Query  727   TGGTAGTCCACGCCCTAAACTATGGATGCTAGTTGTTGCCTTGCTAGTCAAGGCAGTAAT  786
             ||||||||||||||||||||||||||||||||||||||    |||||||    |||||||
Sbjct  706   TGGTAGTCCACGCCCTAAACTATGGATGCTAGTTGTTGGGGAGCTAGTCTCTCCAGTAAT  765

Query  787   GCAGCTAACGCATTAAGCATCCCGCCTGGGGAGTACGGTCGCAAGATTAAAACTCAAAGG  846
             ||||||||||||||||||||||||||||||||||||||||||||||||||||||||||||
Sbjct  766   GCAGCTAACGCATTAAGCATCCCGCCTGGGGAGTACGGTCGCAAGATTAAAACTCAAAGG  825

Query  847   AATAGACGGGGACCCGCACAAGCGGTGGAGCATGTGGTTTAATTCGAAGATACGCGAAGA  906
             ||||||||||||||||||||||||||||||||||||||||||||||||||||| ||||||
Sbjct  826   AATAGACGGGGACCCGCACAAGCGGTGGAGCATGTGGTTTAATTCGAAGATACACGAAGA  885

Query  907   ACCTTACCTAGGCTTGACATTGATAGAATCCGCTAGAGATAGTGGAGTGCCACGCAAGTG  966
             |||||||||||||||||||||||||||||| ||||||||||| |||||||||    | ||
Sbjct  886   ACCTTACCTAGGCTTGACATTGATAGAATCTGCTAGAGATAGCGGAGTGCCAGTTTACTG  945

Query  967   GAGCTTGAAAACAGGTGCTGCACGGCTGTCGTCAGCTCGTGTCGTGAGATGTTGGGTTAA  1026
             ||||||||||||||||||||||||||||||||||||||||||||||||||||||||||||
Sbjct  946   GAGCTTGAAAACAGGTGCTGCACGGCTGTCGTCAGCTCGTGTCGTGAGATGTTGGGTTAA  1005

Query  1027  GTCCCGCAACGAGCGCAACCCTTATCCTTAGTTGCTAGCAGTTAGGCTGAGCACTCTAAG  1086
             ||||||||||||||||||||||   |||||||||||||||||||||||||||||| ||||
Sbjct  1006  GTCCCGCAACGAGCGCAACCCTCGCCCTTAGTTGCTAGCAGTTAGGCTGAGCACTATAAG  1065

Query  1087  GAGACTGCCTTCGTAAGAAGGAGGAAGGCGAGGACGACGTCAAGTCATCATGGCCCTTAC  1146
             | ||||||||| |||||||||||||||| |||||||||||||||||||||||||||||||
Sbjct  1066  GGGACTGCCTTTGTAAGAAGGAGGAAGGTGAGGACGACGTCAAGTCATCATGGCCCTTAC  1125

Query  1147  GCCTAGGGCTACACACGTGCTACAATGGTTGATACAAAGAGATGCAATACTGCGAAGTGG  1206
             |||||||||||||||||||||||||||||||||||||||||| || |||||| |||||||
Sbjct  1126  GCCTAGGGCTACACACGTGCTACAATGGTTGATACAAAGAGAAGCGATACTGTGAAGTGG  1185

Query  1207  AGCCAATCTCAAAAATCAATCTCAGTTCGGATTGTAGGCTGCAACTCGCCTACATGAAGC  1266
               |||||||||||||||||||||||||||||||| |||||||||||||||| ||||||||
Sbjct  1186  GACCAATCTCAAAAATCAATCTCAGTTCGGATTGAAGGCTGCAACTCGCCTTCATGAAGC  1245

Query  1267  TGGAATCGCTAGTAATCGTAAATCAGCAATGTTACGGTGAATACGTTCCCGGGTCTTGTA  1326
             ||||||||||||||||||| ||||||| |||| |||||||||||||||||||||||||||
Sbjct  1246  TGGAATCGCTAGTAATCGTGAATCAGCGATGTCACGGTGAATACGTTCCCGGGTCTTGTA  1305

Query  1327  CTCACCGCCCGTCACACCATGGGAGTTGTATTCGCCTTAAGTCGGAATACTAAATTAGTT  1386
             |||||||||||||||||||||||||||||||||||||||||||||||||||||||||||
Sbjct  1306  CTCACCGCCCGTCACACCATGGGAGTTGTATTCGCCTTAAGTCGGAATACTAAATTAGT-  1364

Query  1387  ACCGC  1391
             |||||
Sbjct  1365  ACCGC  1369
```
