## Supplementary_Data_2 for "State-Threatened Gopher Tortoise (*Gopherus polyphemus*) Gut Microbiome Analysis Reveals Health Insights into Southeastern Florida Population"

```
RID: 8WBGNH9W016
Job Title:seq2192 size=426 counts=76
Program: BLASTN
Database: nt Nucleotide collection (nt)
Query #1: seq2192 size=426 counts=76 Query ID: lcl|Query_7212865 Length: 426

Sequences producing significant alignments:
                                                                  Scientific      Common                     Max    Total Query   E    Per.   Acc.
Description                                                       Name            Name            Taxid      Score  Score cover Value  Ident  Len        Accession
Uncultured bacterium clone B590 16S ribosomal RNA gene, partia... uncultured b... NA              77133      684    684   99%   0.0    95.75  428        MF584337.1
Uncultured bacterium clone 521 16S ribosomal RNA gene, partial... uncultured b... NA              77133      662    662   99%   0.0    94.81  428        OQ027494.1
Mycoplasma sp. B17 chromosome                                     Mycoplasma s... NA              2933905    632    632   100%  2e-176 93.44  656646     CP095758.1
Mycoplasma sp. isolate BAAFS1 chromosome, complete genome         Mycoplasma sp.  NA              2108       627    627   100%  1e-174 93.21  656646     CP091280.1
Uncultured rumen bacterium 16S rRNA, partial sequence, clone:...  uncultured r... NA              136703     621    621   100%  5e-173 92.96  1473       AB615021.1
Uncultured rumen bacterium 16S rRNA, partial sequence, clone:...  uncultured r... NA              136703     621    621   100%  5e-173 92.96  1473       AB614867.1
Uncultured rumen bacterium gene for 16S rRNA, partial sequence... uncultured r... NA              136703     621    621   100%  5e-173 92.96  1474       AB616359.1
Uncultured rumen bacterium gene for 16S rRNA, partial sequence... uncultured r... NA              136703     621    621   100%  5e-173 92.96  1473       AB616265.1
Uncultured rumen bacterium clone L406RT-6-G12 16S ribosomal RN... uncultured r... NA              136703     621    621   100%  5e-173 92.96  1477       GU304581.1
Uncultured bacterium clone 1653 16S ribosomal RNA gene, partia... uncultured b... NA              77133      617    617   93%   7e-172 94.51  401        KP104003.1
Uncultured rumen bacterium clone L406RC-6-D11 16S ribosomal RN... uncultured r... NA              136703     614    614   100%  9e-171 92.49  1476       GU303968.1
Candidatus Mycoplasma girerdii ZYMg-239 gene for 16S rRNA,...     Candidatus M... NA              1318617    610    610   100%  1e-169 92.51  1087       LC554421.1
Candidatus Mycoplasma girerdii ZYMg-227 gene for 16S rRNA,...     Candidatus M... NA              1318617    610    610   100%  1e-169 92.51  1087       LC554420.1
Candidatus Mycoplasma girerdii ZYMg-104 gene for 16S rRNA,...     Candidatus M... NA              1318617    610    610   100%  1e-169 92.51  1087       LC554419.1
Candidatus Mycoplasma girerdii ZYMg-3 gene for 16S rRNA, parti... Candidatus M... NA              1318617    610    610   100%  1e-169 92.51  1087       LC554418.1
Candidatus Malacoplasma girerdii isolate HS41 16S ribosomal RN... Candidatus M... NA              1318617    610    610   100%  1e-169 92.51  710        MF769620.1
Candidatus Mycoplasma girerdii gene for 16S ribosomal RNA,...     Candidatus M... NA              1318617    610    610   100%  1e-169 92.51  1097       LC272066.1
Candidatus Mycoplasma girerdii gene for 16S ribosomal RNA,...     Candidatus M... NA              1318617    610    610   100%  1e-169 92.51  1088       LC272065.1
Mycoplasma sp. HK-4 gene for 16S rRNA, partial sequence           Mycoplasma sp.  NA              2108       610    610   100%  1e-169 92.51  1455       LC777633.1
Candidatus Malacoplasma girerdii isolate UC_B3 chromosome,...     Candidatus M... NA              1318617    610    610   100%  1e-169 92.51  629409     CP020122.1
Uncultured bacterium clone 50 16S ribosomal RNA gene, partial...  uncultured b... NA              77133      610    610   100%  1e-169 92.51  688        KY781879.1
Uncultured bacterium clone 48 16S ribosomal RNA gene, partial...  uncultured b... NA              77133      610    610   100%  1e-169 92.51  692        KY781877.1
Uncultured bacterium clone 43 16S ribosomal RNA gene, partial...  uncultured b... NA              77133      610    610   100%  1e-169 92.51  599        KY781872.1
Uncultured bacterium clone 42 16S ribosomal RNA gene, partial...  uncultured b... NA              77133      610    610   100%  1e-169 92.51  509        KY781871.1
Uncultured bacterium clone 40 16S ribosomal RNA gene, partial...  uncultured b... NA              77133      610    610   100%  1e-169 92.51  690        KY781869.1
Uncultured bacterium clone 39 16S ribosomal RNA gene, partial...  uncultured b... NA              77133      610    610   100%  1e-169 92.51  699        KY781868.1
Uncultured bacterium clone 38 16S ribosomal RNA gene, partial...  uncultured b... NA              77133      610    610   100%  1e-169 92.51  572        KY781867.1
Uncultured bacterium clone 37 16S ribosomal RNA gene, partial...  uncultured b... NA              77133      610    610   100%  1e-169 92.51  608        KY781866.1
Uncultured bacterium clone 36 16S ribosomal RNA gene, partial...  uncultured b... NA              77133      610    610   100%  1e-169 92.51  684        KY781865.1
Uncultured bacterium clone 34 16S ribosomal RNA gene, partial...  uncultured b... NA              77133      610    610   100%  1e-169 92.51  582        KY781863.1
Uncultured bacterium clone 32 16S ribosomal RNA gene, partial...  uncultured b... NA              77133      610    610   100%  1e-169 92.51  688        KY781861.1
Uncultured bacterium clone 31 16S ribosomal RNA gene, partial...  uncultured b... NA              77133      610    610   100%  1e-169 92.51  693        KY781860.1
Uncultured bacterium clone 30 16S ribosomal RNA gene, partial...  uncultured b... NA              77133      610    610   100%  1e-169 92.51  615        KY781859.1
Uncultured bacterium clone 29 16S ribosomal RNA gene, partial...  uncultured b... NA              77133      610    610   100%  1e-169 92.51  675        KY781858.1
Uncultured bacterium clone 28 16S ribosomal RNA gene, partial...  uncultured b... NA              77133      610    610   100%  1e-169 92.51  688        KY781857.1
Uncultured bacterium clone 22 16S ribosomal RNA gene, partial...  uncultured b... NA              77133      610    610   100%  1e-169 92.51  673        KY781851.1
Uncultured bacterium clone 21 16S ribosomal RNA gene, partial...  uncultured b... NA              77133      610    610   100%  1e-169 92.51  651        KY781850.1
Uncultured bacterium clone 20 16S ribosomal RNA gene, partial...  uncultured b... NA              77133      610    610   100%  1e-169 92.51  697        KY781849.1
Uncultured bacterium clone 14 16S ribosomal RNA gene, partial...  uncultured b... NA              77133      610    610   100%  1e-169 92.51  703        KY781843.1
Uncultured bacterium clone 7 16S ribosomal RNA gene, partial...   uncultured b... NA              77133      610    610   100%  1e-169 92.51  688        KY781836.1
Uncultured bacterium clone 6 16S ribosomal RNA gene, partial...   uncultured b... NA              77133      610    610   100%  1e-169 92.51  706        KY781835.1
Uncultured bacterium clone 5 16S ribosomal RNA gene, partial...   uncultured b... NA              77133      610    610   100%  1e-169 92.51  699        KY781834.1
Uncultured bacterium clone 4 16S ribosomal RNA gene, partial...   uncultured b... NA              77133      610    610   100%  1e-169 92.51  698        KY781833.1
Uncultured Mycoplasma sp. clone sq15 16S ribosomal RNA gene,...   uncultured M... NA              167967     610    610   100%  1e-169 92.52  1005       PP396090.1
Candidatus Mycoplasma girerdii strain VCU_M1, complete genome     Candidatus M... NA              1318617    610    610   100%  1e-169 92.51  618983     CP007711.1
Uncultured Mycoplasma sp. partial 16S rRNA gene, clone D04        uncultured M... NA              167967     610    610   100%  1e-169 92.51  1332       HG764211.1
Uncultured Mycoplasma sp. partial 16S rRNA gene, clone D06        uncultured M... NA              167967     610    610   100%  1e-169 92.51  1332       HG764210.1
Uncultured Mycoplasma sp. partial 16S rRNA gene, clone H07        uncultured M... NA              167967     610    610   100%  1e-169 92.51  1332       HG764209.1
Uncultured bacterium clone 22.104-1 16S ribosomal RNA gene,...    uncultured b... NA              77133      610    610   100%  1e-169 92.51  1035       JX871253.1
Uncultured Mycoplasma sp. clone Mnola 16S ribosomal RNA gene,...  uncultured M... NA              167967     610    610   100%  1e-169 92.51  1472       JX508800.1
Uncultured Mycoplasmatales bacterium clone Rc571 16S ribosomal... uncultured M... NA              749450     610    610   100%  1e-169 92.52  1471       JQ617859.1
Uncultured Mycoplasma sp. gene for 16S rRNA, partial sequence,... uncultured M... NA              167967     610    610   100%  1e-169 92.52  1326       AB192228.1
Uncultured Mycoplasma sp. gene for 16S rRNA, partial sequence,... uncultured M... NA              167967     610    610   100%  1e-169 92.52  1326       AB192181.1
Uncultured Mycoplasma sp. gene for 16S rRNA, partial sequence,... uncultured M... NA              167967     610    610   100%  1e-169 92.49  1430       AB089057.1
Uncultured Mycoplasma sp. gene for 16S rRNA, partial sequence,... uncultured M... NA              167967     610    610   100%  1e-169 92.52  1326       AB089053.1
Uncultured bacterium clone 24 16S ribosomal RNA gene, partial...  uncultured b... NA              77133      606    606   100%  1e-168 92.27  596        KY781853.1
Uncultured Mycoplasma sp. clone abc370 16S ribosomal RNA gene,... uncultured M... NA              167967     606    606   99%   1e-168 92.47  428        MN665175.1
Uncultured Mycoplasma sp. partial 16S rRNA gene, clone D08        uncultured M... NA              167967     604    604   100%  5e-168 92.27  1332       HG764212.1
Uncultured bacterium clone 41 16S ribosomal RNA gene, partial...  uncultured b... NA              77133      603    603   100%  2e-167 92.27  690        KY781870.1
Uncultured bacterium clone 3 16S ribosomal RNA gene, partial...   uncultured b... NA              77133      603    603   100%  2e-167 92.27  613        KY781832.1
Uncultured bacterium clone 2 16S ribosomal RNA gene, partial...   uncultured b... NA              77133      603    603   100%  2e-167 92.27  690        KY781831.1
Uncultured bacterium clone 56 16S ribosomal RNA gene, partial...  uncultured b... NA              77133      599    599   100%  2e-166 92.04  683        KY781884.1
Uncultured bacterium clone 45 16S ribosomal RNA gene, partial...  uncultured b... NA              77133      599    599   100%  2e-166 92.04  693        KY781874.1
Uncultured bacterium clone 8 16S ribosomal RNA gene, partial...   uncultured b... NA              77133      595    595   100%  3e-165 91.57  684        KY781837.1
Uncultured bacterium clone 15 16S ribosomal RNA gene, partial...  uncultured b... NA              77133      588    588   100%  5e-163 91.10  577        KY781844.1
Uncultured rumen bacterium 16S rRNA, partial sequence, clone:...  uncultured r... NA              136703     588    588   100%  5e-163 91.55  1474       AB555181.1
Uncultured bacterium clone B1369 16S ribosomal RNA gene, parti... uncultured b... NA              77133      579    579   99%   3e-160 91.29  429        MF585105.1
Uncultured Firmicutes bacterium clone Md131_11 16S ribosomal R... uncultured F... NA              344338     577    577   100%  1e-159 91.08  1474       KM650840.1
Uncultured Mollicutes bacterium clone RsaTcA_113 16S ribosomal... uncultured M... NA              220137     577    577   100%  1e-159 91.14  1432       JQ993512.1
Uncultured Firmicutes bacterium clone Cf4-93 16S ribosomal RNA... uncultured F... NA              344338     571    571   100%  5e-158 90.85  1392       GQ502567.1
Mycoplasma sp. 16S rRNA gene, partial                             Mycoplasma sp.  NA              2108       571    571   99%   5e-158 91.04  1147       AJ132469.1
Uncultured Firmicutes bacterium clone Md131_7 16S ribosomal RN... uncultured F... NA              344338     566    566   100%  3e-156 90.61  1452       KM650864.1
Uncultured bacterium clone MgKI1c001F07 16S ribosomal RNA gene... uncultured b... NA              77133      566    566   100%  3e-156 90.61  1479       KP690933.1
Uncultured bacterium clone B2507 16S ribosomal RNA gene, parti... uncultured b... NA              77133      562    562   99%   3e-155 90.63  429        MF586209.1
Uncultured bacterium clone 18 16S ribosomal RNA gene, partial...  uncultured b... NA              77133      560    560   93%   1e-154 92.00  649        KY781847.1
Uncultured Mollicutes bacterium gene for 16S rRNA, partial...     uncultured M... NA              220137     560    560   100%  1e-154 90.42  1330       LC198349.1
Uncultured bacterium clone C50 16S ribosomal RNA gene, partial... uncultured b... NA              77133      555    555   100%  5e-153 90.14  520        HQ180159.1
Uncultured Firmicutes bacterium clone Cf4-19 16S ribosomal RNA... uncultured F... NA              344338     555    555   100%  5e-153 90.14  1395       GQ502581.1
Uncultured Mollicutes bacterium clone HsjTcC_106 16S ribosomal... uncultured M... NA              220137     549    549   100%  3e-151 89.91  1432       JQ993492.1
Uncultured Mollicutes bacterium clone HsjTcB_50 16S ribosomal...  uncultured M... NA              220137     549    549   100%  3e-151 89.91  1432       JQ993479.1
Uncultured Firmicutes bacterium clone Cf4-65 16S ribosomal RNA... uncultured F... NA              344338     544    544   100%  1e-149 89.70  1381       GQ502548.1
Uncultured bacterium clone WA_aaa03d11 16S ribosomal RNA gene,... uncultured b... NA              77133      538    538   100%  6e-148 89.49  1371       EU473607.1
Uncultured bacterium clone WA_aaa01a03 16S ribosomal RNA gene,... uncultured b... NA              77133      538    538   100%  6e-148 89.49  1292       EU473533.1
Uncultured bacterium clone AE2_aaa03d07 16S ribosomal RNA gene... uncultured b... NA              77133      538    538   100%  6e-148 89.53  1371       EU471802.1
Uncultured Mycoplasma sp. gene for 16S rRNA, partial sequence,... uncultured M... NA              167967     538    538   100%  6e-148 89.46  1327       AB299549.1
Uncultured bacterium partial 16S rRNA gene, clone CbDvB59         uncultured b... NA              77133      536    536   100%  2e-147 89.46  1469       FN377796.1
Uncultured bacterium clone MgKI1c002B12 16S ribosomal RNA gene... uncultured b... NA              77133      532    532   100%  3e-146 89.20  1481       KP690955.1
Uncultured bacterium gene for 16S ribosomal RNA, partial...       uncultured b... NA              77133      532    532   100%  3e-146 89.30  1474       AB893931.1
Uncultured bacterium clone WA_aaa04g10 16S ribosomal RNA gene,... uncultured b... NA              77133      532    532   100%  3e-146 89.25  1393       EU779362.1
Uncultured Ureaplasma sp. clone OTU_1 16S ribosomal RNA gene,...  uncultured U... NA              293430     529    529   99%   3e-145 89.25  429        MN955326.1
Uncultured bacterium gene for 16S rRNA, partial sequence, clon... uncultured b... NA              77133      527    527   100%  1e-144 88.99  1371       AB062770.1
Uncultured Mycoplasma sp. clone K.b-1 16S ribosomal RNA gene,...  uncultured M... NA              167967     523    523   100%  2e-143 88.84  907        HM031446.1
Uncultured rumen bacterium clone L406RC-6-D12 16S ribosomal RN... uncultured r... NA              136703     521    521   88%   6e-143 91.29  720        GU303969.1
Uncultured Ureaplasma sp. clone LP1-P10 16S ribosomal RNA gene... uncultured U... NA              293430     518    518   100%  7e-142 88.60  575        JX453985.1
Uncultured Ureaplasma sp. clone LP1-C02 16S ribosomal RNA gene... uncultured U... NA              293430     518    518   100%  7e-142 88.60  609        JX453794.1
Uncultured Mycoplasmataceae bacterium clone LP1-B08 16S...        uncultured M... NA              300027     518    518   100%  7e-142 88.60  564        JX453782.1
Uncultured Mycoplasma sp. clone C10 16S ribosomal RNA gene,...    uncultured M... NA              167967     518    518   100%  7e-142 88.60  1456       DQ340196.1
Uncultured bacterium clone AE2_aaa03d08 16S ribosomal RNA gene... uncultured b... NA              77133      516    516   100%  3e-141 88.55  1373       EU471695.1
Uncultured bacterium clone horsej_aai91e10 16S ribosomal RNA...   uncultured b... NA              77133      516    516   100%  3e-141 88.55  1380       EU463716.1
Uncultured bacterium clone WDol03Rect1A08 16S ribosomal RNA...    uncultured b... NA              77133      512    512   100%  3e-140 88.37  761        KC257899.1


Alignments:

>Uncultured bacterium clone B590 16S ribosomal RNA gene, partial sequence
Sequence ID: MF584337.1 Length: 428
Range 1: 1 to 424

Score:684 bits(370), Expect:0.0,
Identities:406/424(96%),  Gaps:0/424(0%), Strand: Plus/Plus

Query  3    TAGGGAATTTTTCACAATGGGCGAAAGCCTGATGGAGCAATACCGCGTGGATGATGAAGG  62
            ||||||||||||||||||||||||||||||||||||||||||||||||||||||||||||
Sbjct  1    TAGGGAATTTTTCACAATGGGCGAAAGCCTGATGGAGCAATACCGCGTGGATGATGAAGG  60

Query  63   TCTTCGGATCGTAAAATCCTTTTATAAGGGACGAATGATATAAGTAGGAAATGATTTATA  122
            |||| |||| ||||||||||||||| |||||||||||| ||||||||||||||||||||
Sbjct  61   TCTTTGGATTGTAAAATCCTTTTATTAGGGACGAATGACATAAGTAGGAAATGATTTATG  120

Query  123  TTTGACTGTACCTTTTGAATAAGTAACGGCAAACTATGTGCCAGCAGCCGCGGTAATACA  182
            ||||||||||||||||||||||||||||||||||||||||||||||||||||||||||||
Sbjct  121  TTTGACTGTACCTTTTGAATAAGTAACGGCAAACTATGTGCCAGCAGCCGCGGTAATACA  180

Query  183  TAGGTTACAAGCGTTATCCGGATTTACTGGGCGTAAAGCGAGCGCAGGCTGGTTTATAAG  242
            ||||||||||||||||||||||||||||||||||||||||||||||||||| |||| |||
Sbjct  181  TAGGTTACAAGCGTTATCCGGATTTACTGGGCGTAAAGCGAGCGCAGGCTGATTTACAAG  240

Query  243  TCTAGTGTTAAATACGATTGCTTAACAATCGTTTGCATTGGAAACTATAAACCTAGAGTG  302
            ||| |||||||||||  ||||| |||||  || ||||||||||||| |||  ||||||||
Sbjct  241  TCTGGTGTTAAATACAGTTGCTCAACAACTGTATGCATTGGAAACTGTAAGTCTAGAGTG  300

Query  303  TGATAGGGAGTTCTGGAACTCCATGTGGAGCGGTGGAATGCGTAGATATATGGAAGAACA  362
            ||||||||||||||||||||||||||||||||||||||||||||||||||||||||||||
Sbjct  301  TGATAGGGAGTTCTGGAACTCCATGTGGAGCGGTGGAATGCGTAGATATATGGAAGAACA  360

Query  363  CCAGTGGCGAAAGCGAGAACTTAGGTCACTACTGACGCTTAGGCTCGAAAGTGTGGGGAG  422
            ||||||||||||||||||||||||||||| ||||||||||||||||||||||||||||||
Sbjct  361  CCAGTGGCGAAAGCGAGAACTTAGGTCACAACTGACGCTTAGGCTCGAAAGTGTGGGGAG  420

Query  423  CAAA  426
            ||||
Sbjct  421  CAAA  424


>Uncultured bacterium clone 521 16S ribosomal RNA gene, partial sequence
Sequence ID: OQ027494.1 Length: 428
Range 1: 1 to 424

Score:662 bits(358), Expect:0.0,
Identities:402/424(95%),  Gaps:0/424(0%), Strand: Plus/Plus

Query  3    TAGGGAATTTTTCACAATGGGCGAAAGCCTGATGGAGCAATACCGCGTGGATGATGAAGG  62
            ||||||||||||||||||||||||||||||||||||||||||||||||||||||||||||
Sbjct  1    TAGGGAATTTTTCACAATGGGCGAAAGCCTGATGGAGCAATACCGCGTGGATGATGAAGG  60

Query  63   TCTTCGGATCGTAAAATCCTTTTATAAGGGACGAATGATATAAGTAGGAAATGATTTATA  122
            |||| |||| ||||||||||||||| |||||||||||| | |||||||||||||||| |
Sbjct  61   TCTTTGGATTGTAAAATCCTTTTATTAGGGACGAATGACACAAGTAGGAAATGATTTGTG  120

Query  123  TTTGACTGTACCTTTTGAATAAGTAACGGCAAACTATGTGCCAGCAGCCGCGGTAATACA  182
            ||||||||||||||||||||||||||||||||||||||||||||||||||||||||||||
Sbjct  121  TTTGACTGTACCTTTTGAATAAGTAACGGCAAACTATGTGCCAGCAGCCGCGGTAATACA  180

Query  183  TAGGTTACAAGCGTTATCCGGATTTACTGGGCGTAAAGCGAGCGCAGGCTGGTTTATAAG  242
            ||||||||||||||||||||||||||||||||||||||||||||||||||| || | |||
Sbjct  181  TAGGTTACAAGCGTTATCCGGATTTACTGGGCGTAAAGCGAGCGCAGGCTGATTCACAAG  240

Query  243  TCTAGTGTTAAATACGATTGCTTAACAATCGTTTGCATTGGAAACTATAAACCTAGAGTG  302
            ||| ||||||||| |  ||||| |||||  |||||||||||||||| | |  ||||||||
Sbjct  241  TCTGGTGTTAAATGCAGTTGCTCAACAACTGTTTGCATTGGAAACTGTGAGTCTAGAGTG  300

Query  303  TGATAGGGAGTTCTGGAACTCCATGTGGAGCGGTGGAATGCGTAGATATATGGAAGAACA  362
            ||||||||||||||||||||||||||||||||||||||||||||||||||||||||||||
Sbjct  301  TGATAGGGAGTTCTGGAACTCCATGTGGAGCGGTGGAATGCGTAGATATATGGAAGAACA  360

Query  363  CCAGTGGCGAAAGCGAGAACTTAGGTCACTACTGACGCTTAGGCTCGAAAGTGTGGGGAG  422
            ||||||||||||||||||||||||||||| ||||||||||||||||||||||||||||||
Sbjct  361  CCAGTGGCGAAAGCGAGAACTTAGGTCACAACTGACGCTTAGGCTCGAAAGTGTGGGGAG  420

Query  423  CAAA  426
            ||||
Sbjct  421  CAAA  424


>MAG: Mycoplasma sp. B17 chromosome
Sequence ID: CP095758.1 Length: 656646
Range 1: 277852 to 278277

Score:632 bits(342), Expect:2e-176,
Identities:399/427(93%),  Gaps:2/427(0%), Strand: Plus/Minus

Query  1       AGTAGGGAATTTTTCACAATGGGCGAAAGCCTGATGGAGCAATACCGCGTGGATGATGAA  60
               ||||||||||||||||||||||||||||||||||||||||||||||||||||||||||||
Sbjct  278277  AGTAGGGAATTTTTCACAATGGGCGAAAGCCTGATGGAGCAATACCGCGTGGATGATGAA  278218

Query  61      GGTCTTCGGATCGTAAAATCCTTTTATAAGGGACGAATGATA-TAAGTAGGAAATGATTT  119
               ||||||||||| ||||||||||||||| |||||||||||  | |||| |||||||| ||
Sbjct  278217  GGTCTTCGGATTGTAAAATCCTTTTATTAGGGACGAATGGCACTAAG-AGGAAATGCTTA  278159

Query  120     ATATTTGACTGTACCTTTTGAATAAGTAACGGCAAACTATGTGCCAGCAGCCGCGGTAAT  179
                |  ||||||||||||||||||||||||||||||||||||||||||||||||||||||||
Sbjct  278158  GTGATTGACTGTACCTTTTGAATAAGTAACGGCAAACTATGTGCCAGCAGCCGCGGTAAT  278099

Query  180     ACATAGGTTACAAGCGTTATCCGGATTTACTGGGCGTAAAGCGAGCGCAGGCTGGTTTAT  239
               |||||||||||||||||||||||||||||||||||||||||||||||| ||||| | ||
Sbjct  278098  ACATAGGTTACAAGCGTTATCCGGATTTACTGGGCGTAAAGCGAGCGCGGGCTGATATAC  278039

Query  240     AAGTCTAGTGTTAAATACGATTGCTTAACAATCGTTTGCATTGGAAACTATAAACCTAGA  299
               ||||||||||| |||| |  ||||| |||||  |||||||||||||||| || | |||||
Sbjct  278038  AAGTCTAGTGTGAAATGCAGTTGCTCAACAACTGTTTGCATTGGAAACTGTACATCTAGA  277979

Query  300     GTGTGATAGGGAGTTCTGGAACTCCATGTGGAGCGGTGGAATGCGTAGATATATGGAAGA  359
               ||||||||||||||||||||||||||| ||||||||||||||||||||||||||||||||
Sbjct  277978  GTGTGATAGGGAGTTCTGGAACTCCATATGGAGCGGTGGAATGCGTAGATATATGGAAGA  277919

Query  360     ACACCAGTGGCGAAAGCGAGAACTTAGGTCACTACTGACGCTTAGGCTCGAAAGTGTGGG  419
               ||||| |||||||||||||||||||||||||| |||||||||||||||||||||||||||
Sbjct  277918  ACACCTGTGGCGAAAGCGAGAACTTAGGTCACAACTGACGCTTAGGCTCGAAAGTGTGGG  277859

Query  420     GAGCAAA  426
               |||||||
Sbjct  277858  GAGCAAA  277852


>MAG: Mycoplasma sp. isolate BAAFS1 chromosome, complete genome
Sequence ID: CP091280.1 Length: 656646
Range 1: 277852 to 278277

Score:627 bits(339), Expect:1e-174,
Identities:398/427(93%),  Gaps:2/427(0%), Strand: Plus/Minus

Query  1       AGTAGGGAATTTTTCACAATGGGCGAAAGCCTGATGGAGCAATACCGCGTGGATGATGAA  60
               ||||||||||||||||||||||||||||||||||||||||||||||||||||||||||||
Sbjct  278277  AGTAGGGAATTTTTCACAATGGGCGAAAGCCTGATGGAGCAATACCGCGTGGATGATGAA  278218

Query  61      GGTCTTCGGATCGTAAAATCCTTTTATAAGGGACGAATGATA-TAAGTAGGAAATGATTT  119
               ||||||||||| ||||||||||||||| |||||||||||  | |||| |||||||| ||
Sbjct  278217  GGTCTTCGGATTGTAAAATCCTTTTATTAGGGACGAATGGCACTAAG-AGGAAATGCTTA  278159

Query  120     ATATTTGACTGTACCTTTTGAATAAGTAACGGCAAACTATGTGCCAGCAGCCGCGGTAAT  179
                |  ||||||||||||||||||||||||||||||||||||||||||||||||||||||||
Sbjct  278158  GTGATTGACTGTACCTTTTGAATAAGTAACGGCAAACTATGTGCCAGCAGCCGCGGTAAT  278099

Query  180     ACATAGGTTACAAGCGTTATCCGGATTTACTGGGCGTAAAGCGAGCGCAGGCTGGTTTAT  239
               |||||||||||||||||||||||||||||||||||||||||||||||| ||||| | ||
Sbjct  278098  ACATAGGTTACAAGCGTTATCCGGATTTACTGGGCGTAAAGCGAGCGCGGGCTGATATAC  278039

Query  240     AAGTCTAGTGTTAAATACGATTGCTTAACAATCGTTTGCATTGGAAACTATAAACCTAGA  299
               ||||||||||| |||| |  ||||| |||||  |||||||||||||||| || | |||||
Sbjct  278038  AAGTCTAGTGTAAAATGCAGTTGCTCAACAACTGTTTGCATTGGAAACTGTACATCTAGA  277979

Query  300     GTGTGATAGGGAGTTCTGGAACTCCATGTGGAGCGGTGGAATGCGTAGATATATGGAAGA  359
               ||||||||||||||||||||||||||| ||||||||||||||||||||||||||||||||
Sbjct  277978  GTGTGATAGGGAGTTCTGGAACTCCATATGGAGCGGTGGAATGCGTAGATATATGGAAGA  277919

Query  360     ACACCAGTGGCGAAAGCGAGAACTTAGGTCACTACTGACGCTTAGGCTCGAAAGTGTGGG  419
               ||||| ||||||||||||||||||||| |||| |||||||||||||||||||||||||||
Sbjct  277918  ACACCTGTGGCGAAAGCGAGAACTTAGTTCACAACTGACGCTTAGGCTCGAAAGTGTGGG  277859

Query  420     GAGCAAA  426
               |||||||
Sbjct  277858  GAGCAAA  277852


>Uncultured rumen bacterium 16S rRNA, partial sequence, clone: R-B-CA197
Sequence ID: AB615021.1 Length: 1473
Range 1: 341 to 766

Score:621 bits(336), Expect:5e-173,
Identities:396/426(93%),  Gaps:0/426(0%), Strand: Plus/Plus

Query  1    AGTAGGGAATTTTTCACAATGGGCGAAAGCCTGATGGAGCAATACCGCGTGGATGATGAA  60
            ||||||||||||||||||||||||||||||||||||||||||||||||||| ||||||||
Sbjct  341  AGTAGGGAATTTTTCACAATGGGCGAAAGCCTGATGGAGCAATACCGCGTGAATGATGAA  400

Query  61   GGTCTTCGGATCGTAAAATCCTTTTATAAGGGACGAATGATATAAGTAGGAAATGATTTA  120
            ||||||||||| ||||||| ||||||||||||||||| ||| | |  |||||||| ||
Sbjct  401  GGTCTTCGGATTGTAAAATTCTTTTATAAGGGACGAACGATGTGAACAGGAAATGGTTCG  460

Query  121  TATTTGACTGTACCTTTTGAATAAGTAACGGCAAACTATGTGCCAGCAGCCGCGGTAATA  180
             |  ||||||||||||||||||||||||||||||||||||||||||||||||||||||||
Sbjct  461  CAAGTGACTGTACCTTTTGAATAAGTAACGGCAAACTATGTGCCAGCAGCCGCGGTAATA  520

Query  181  CATAGGTTACAAGCGTTATCCGGATTTACTGGGCGTAAAGCGAGCGCAGGCTGGTTTATA  240
            ||||||||||||||||||||||||||||||||||||||||||||||||||||| |||| |
Sbjct  521  CATAGGTTACAAGCGTTATCCGGATTTACTGGGCGTAAAGCGAGCGCAGGCTGATTTACA  580

Query  241  AGTCTAGTGTTAAATACGATTGCTTAACAATCGTTTGCATTGGAAACTATAAACCTAGAG  300
            ||||| ||||||||| |  |||||||||||  || ||||||||||||| |||| ||||||
Sbjct  581  AGTCTGGTGTTAAATGCATTTGCTTAACAAATGTATGCATTGGAAACTGTAAATCTAGAG  640

Query  301  TGTGATAGGGAGTTCTGGAACTCCATGTGGAGCGGTGGAATGCGTAGATATATGGAAGAA  360
            |||||||||||||||||||||||||| ||||||||||||||| |||||||||||||||||
Sbjct  641  TGTGATAGGGAGTTCTGGAACTCCATATGGAGCGGTGGAATGTGTAGATATATGGAAGAA  700

Query  361  CACCAGTGGCGAAAGCGAGAACTTAGGTCACTACTGACGCTTAGGCTCGAAAGTGTGGGG  420
            |||| |||||||||||||||||||||||||| ||||||||||||||||||||||||||||
Sbjct  701  CACCTGTGGCGAAAGCGAGAACTTAGGTCACAACTGACGCTTAGGCTCGAAAGTGTGGGG  760

Query  421  AGCAAA  426
            | ||||
Sbjct  761  ATCAAA  766
```
