## Supplementary figures and images for "State-Threatened Gopher Tortoise (*Gopherus polyphemus*) Gut Microbiome Analysis Reveals Health Insights into Southeastern Florida Population"

### Supplementary_Figure_1

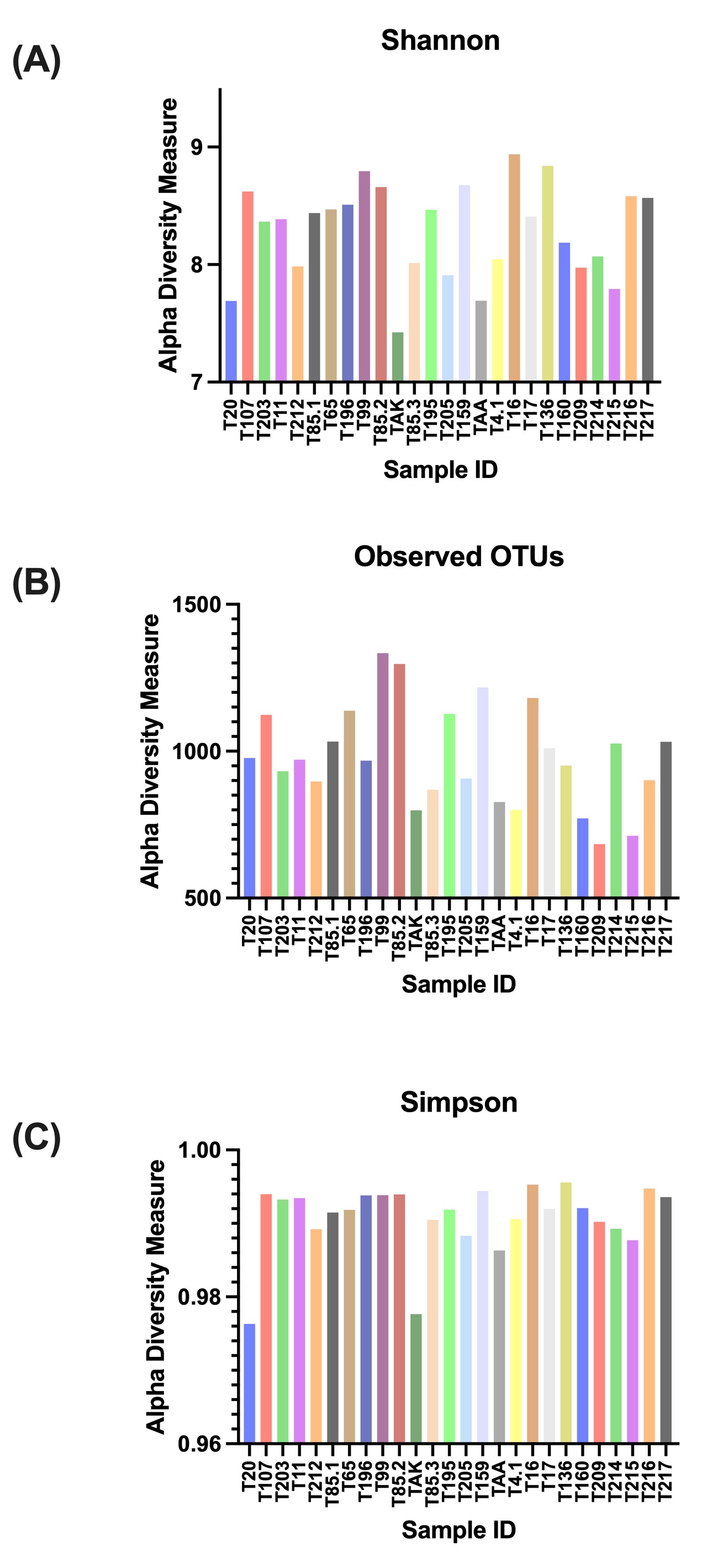

### Supplementary_Figure_2

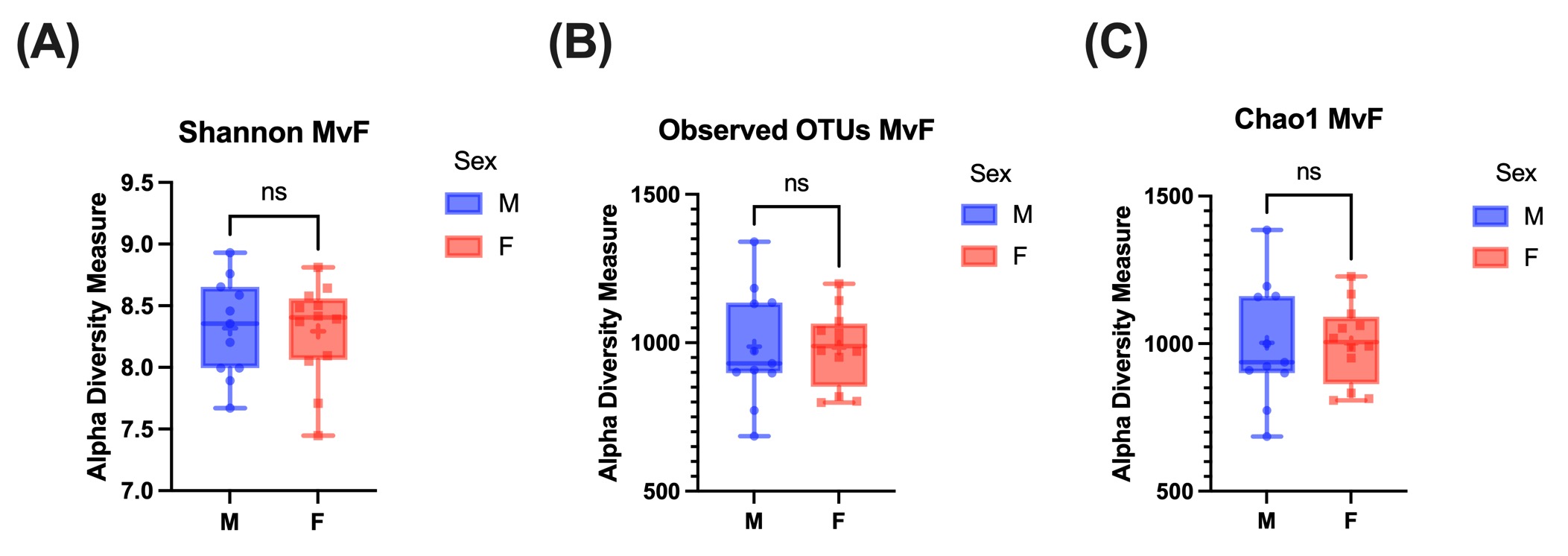

### Supplementary_Figure_3

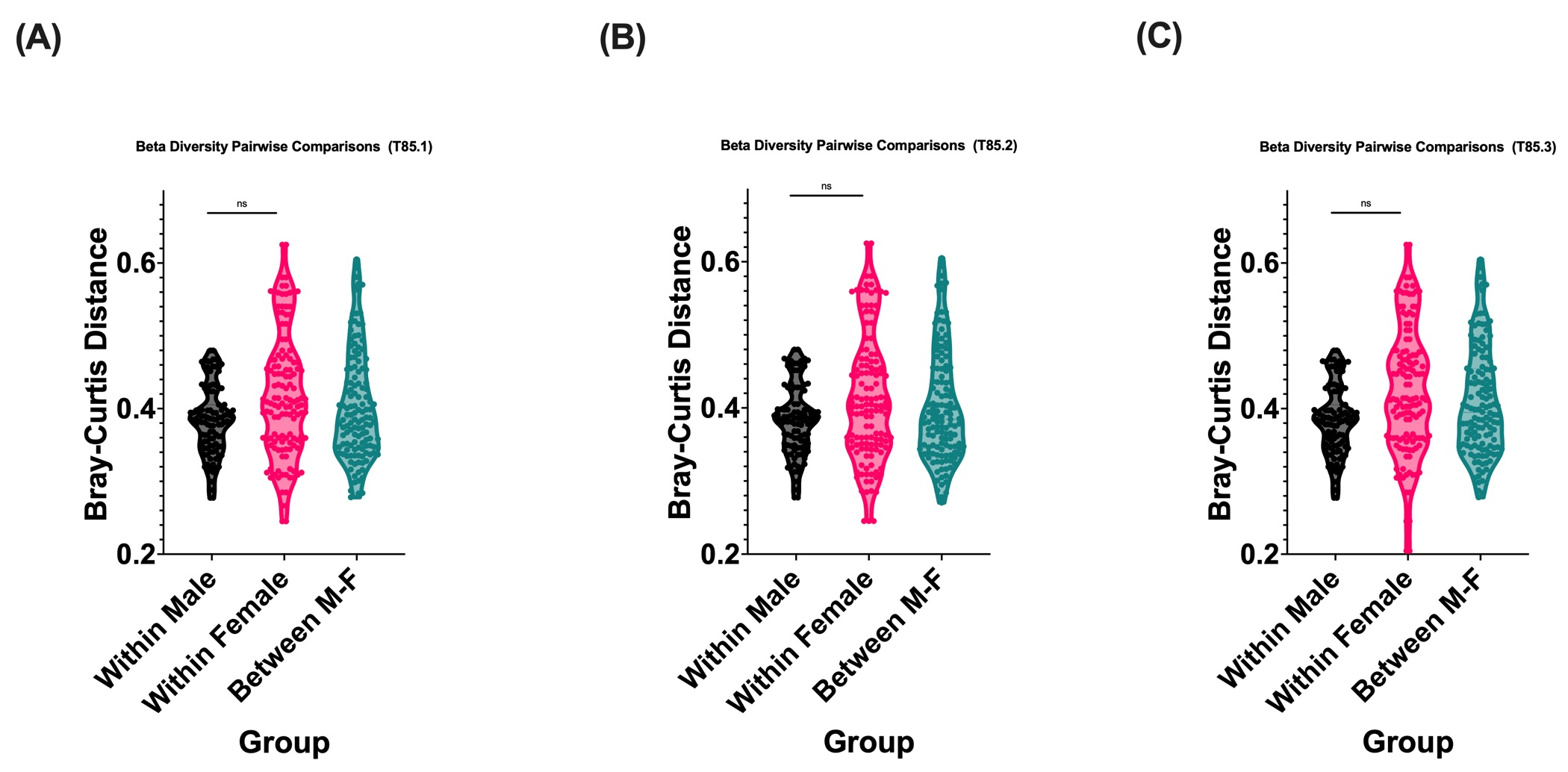
